# Single-Nucleus Profiling Reveals Neuronal Metabolic Collapse During the Transition to Alzheimer’s Disease Dementia

**DOI:** 10.64898/2026.10.26.761376

**Authors:** Songjian Lu, Nitesh Enduru, Citu Citu, Chia-Hao Tung, Zhongming Zhao

**Author notes:** Correspondence should be addressed to: Zhongming Zhao, Ph.D., Center for Precision Health, McWilliams School of Biomedical Informatics, The University of Texas Health Science Center at Houston, 7000 Fannin St Suite 600, Houston, Texas 77030.

## Abstract

Current Alzheimer’s disease (AD) neuropathological metrics, including Thal phasing, and Braak staging, often do not accurately predict cognitive status: many cognitively resilient individuals have substantial pathological burdens, whereas some patients with dementia show minimal classical AD pathology. Here, we addressed this discrepancy by investigating the molecular mechanisms underlying AD dementia using single-nucleus RNA-sequencing and single-nucleus multiome datasets. To eliminate cross-modality scaling artifacts and compositional biases inherent in neurodegenerative tissues, we normalized the datasets using our novel framework, bypassing scaling issues of conventional methods that may obscure critical biological signals of interest. We identified neuron-specific suppression of nuclear-encoded mitochondrial genes and their upstream mitochondrial biogenesis signaling (MBS) as primary drivers of clinical dementia. By integrating multi-omic quantitative trait loci, genome-wide association studies, and epigenomic profiles, we delineated genetic and epigenetic changes underlying MBS suppression. This study provides a systematic genomic framework for understanding the mechanisms of cognitive resilience.

---

Alzheimer’s disease (AD) is the most prevalent neurodegenerative disorder among older adults^1, 2^, and is characterized clinically by progressive declines in memory, executive function, and overall cognitive performance; collectively these measures are referred to as AD dementia. Traditionally, AD diagnosis and staging have relied on postmortem neuropathological assessments. Gold standard measures include Thal phases for amyloid-beta plaque distribution, Braak stages for tau neurofibrillary tangle spread, and Consortium to Establish a Registry for Alzheimer’s Disease (CERAD) scores for neurotic plaque density^3^. Recently, the Seattle Alzheimer’s Disease Brain Cell Atlas (SEA-AD) cohort introduced an overall AD Neuropathological Change (ADNC) score^4^ to provide a more integrated measure of disease severity. However, these pathological markers often fail to predict clinical status. A significant subset of “resilient” individuals maintains cognitive health despite high pathological scores, while others suffer from dementia despite relatively low scores^4–6^. This clinical-pathological mismatch suggests that traditional scores alone are insufficient to define the molecular threshold of dementia.

To identify molecular drivers of cognitive decline, research has shifted from bulk^7–9^ to single-cell transcriptomics^10–12^. Although initial bulk RNA-seq studies mapped broad gene expression changes in AD brains^7–9^, varying cellular compositions confounded their findings. To overcome this, recent single-nucleus RNA-seq (snRNA-seq) ^6, 13^ and multiome (snMultiome; paired snRNA-seq/snATAC-seq)^4, 14–16^ datasets provide fine-grained, cell-type-specific resolution of AD signatures. Notably, the SEA-AD integrated multimodal profiling with quantitative neuropathology^4^, mapping clinical trajectories against molecular states to better distinguish the pathological burden from symptomatic progression.

Leveraging the SEA-AD cohort, in this study we mapped the specific molecular programs that underpin the transition to AD dementia. To overcome technical noise across modalities, we applied **C**ount based on **L**inearized **T**ranscriptome Size (**CLTS**)^17^, a normalization framework designed to eliminate scaling artifacts snRNA-seq data, and extended it to snATAC-seq data. By applying CLTS, we resolved crucial biological signals that remain obscured under conventional normalization approaches, such as Counts Per 10 Thousand (CP10K) and Transcripts Per Million (TPM) for transcriptomics or Term Frequency–Inverse Document Frequency (TF-IDF)^18, 19^ for chromatin accessibility. Integrating these normalized snRNA-seq and snATAC-seq profiles with xQTL mapping enabled us to delineate the genetic and epigenetic architectures repressing metabolic function in dementia. Specifically, we report that the downstream suppression of nuclear-encoded mitochondrial genes (NEMGs) and their upstream mitochondrial biogenesis pathways in neurons is a defining feature of AD dementia progression.

## Results

### Neuropathological profiling and clinical staging of the SEA-AD cohort

The SEA-AD cohort comprised 84 subjects, including 9 normal controls (No-AD) and 75 AD-affected individuals (42 demented [AD-dem] and 33 non-demented [AD-non-dem])^4^. This separation provides an opportunity to investigate the molecular drivers of cognitive decline and dementia from general AD pathology (**Extended Data Fig. 1a**).

Standard neuropathological markers (Thal phase, Braak stage, CERAD, and ADNC) are insufficient to reliably predict clinical cognitive status (**Extended Data Fig. 1b-e**). Notably, over 40% of the AD-non-dem samples exhibited advanced pathology (Thal 4–5, Braak V–VI, Moderate-to-Frequent CERAD scores, and Intermediate-High ADNC score), suggesting a high degree of cognitive resilience. Conversely, a substantial portion of the AD-dem group exhibited relatively low pathological scores (Thal 1–3, Braak I–IV, Absent-to-Sparse CERAD scores, and Low ADNC scores).

To address the gap in how pathology translates to clinical symptoms, this study identified cellular and molecular drivers of AD dementia using snRNA-seq and snMultiome data from the SEA-AD middle temporal gyrus (MTG)^4^. We validated these findings across additional SEA-AD brain regions and in independent cohorts^6^. Crucially, donor age and post-mortem interval (PMI) did not differ significantly across the clinical groups (**Extended Data Fig. 1f, g**), confirming that our differential expression reflects true disease biology rather than demographic confounders. The analytical pipeline is shown on **Fig. 1**.

**Fig. 1.**
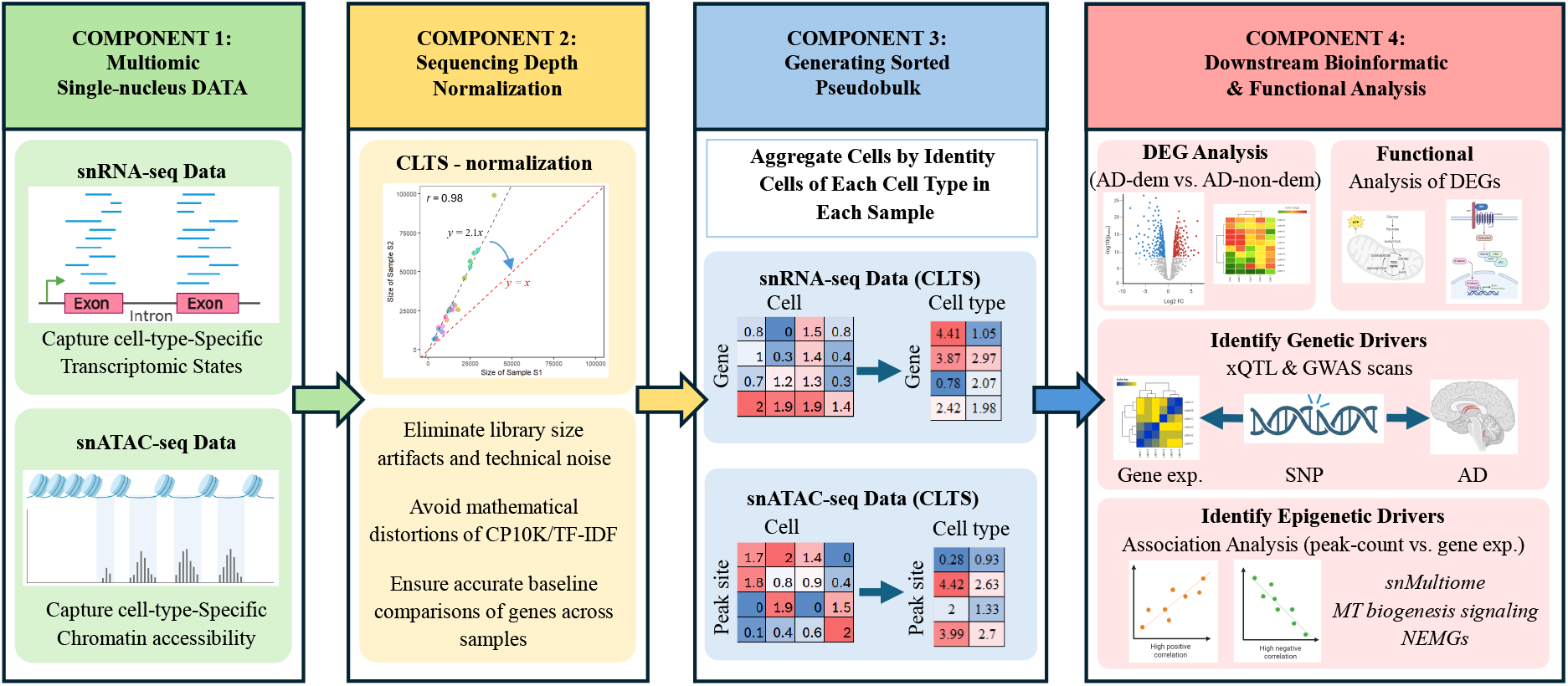
Overview of the analytical framework. Schematic workflow illustrating single-nucleus RNA sequencing (snRNA-seq) and single-nucleus ATAC sequencing (snATAC-seq) data processing. Datasets were normalized using CLTS-mtx, a new normalization method without scaling artifacts, followed by the generation of cell type–specific pseudobulk RNA-seq profiles for each sample. These profiles were used to evaluate functional gene expression changes and epigenetic alterations associated with Alzheimer’s disease (AD) dementia. (Icons created with BioRender.com).

### Normalization of snRNA-seq and snATAC-seq data using CLTS

Data generated from sequencing-based technologies, such as snRNA-seq and snATAC-seq data, are inherently sensitive to sequencing depth. Consequently, the raw count profiles of cells cannot be directly compared across samples. For instance, in the SEA-AD MTG snRNA-seq dataset, raw count expressions of *ST6GALNAC3* in L5 ET and microglia cells exhibit significant artificial variance between samples S1 and S2 (**Fig. 2a**, **raw counts**), leading to a biased comparison if left uncorrected.

**Fig. 2.**
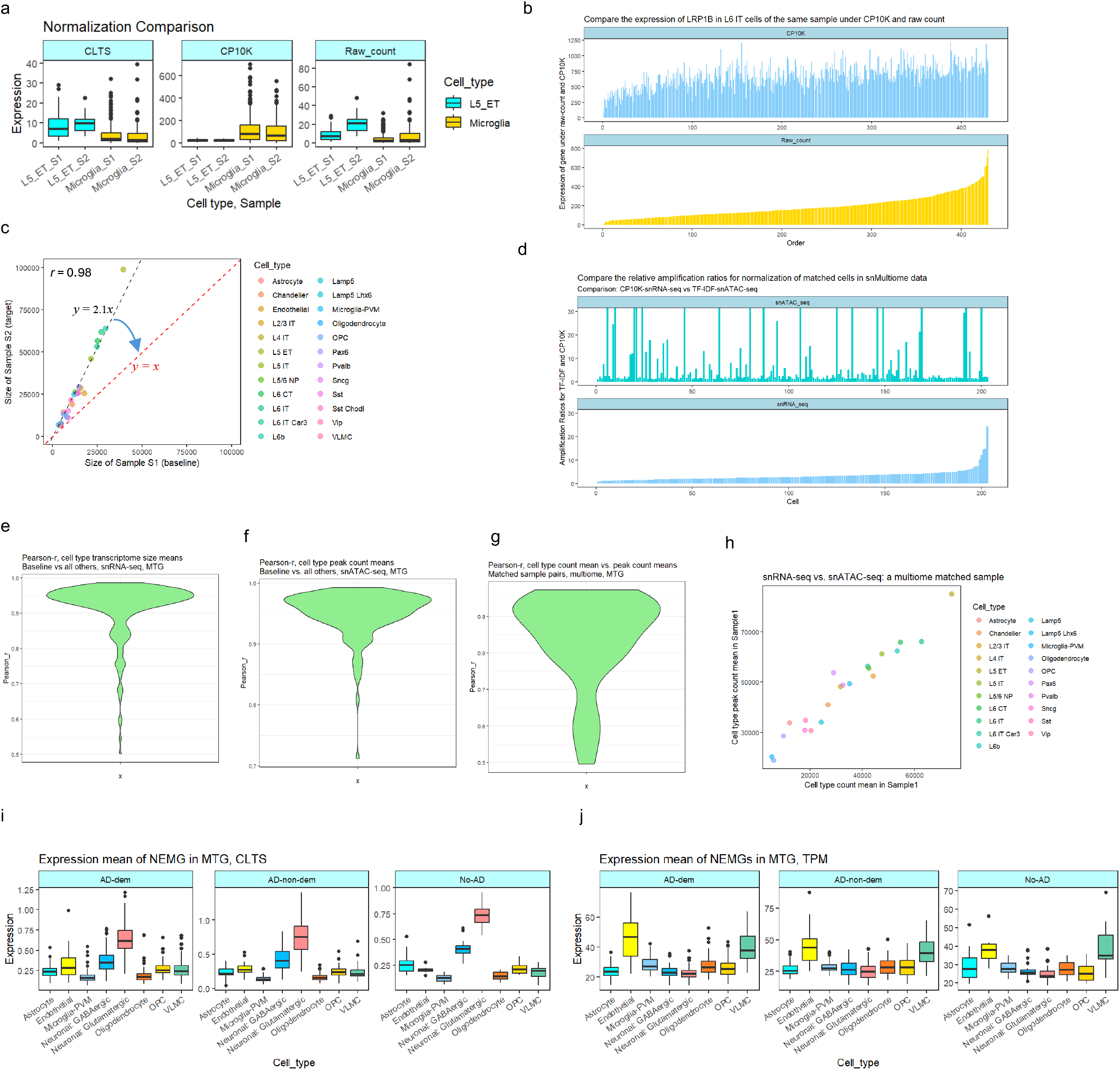
Benchmarking CLTS Normalization for Cross-Sample and Cross-Modality Data Integration. **a**, Comparative expression of *ST6GALNAC3* in microglia and L5 ET neurons across samples S1 and S2, contrasting raw counts with CP10K and CLTS normalization. **b**, Expressions of *LRP1B* in L6 IT cells (sample S1) under raw count and standard CP10K normalization. Cells were sorted by their raw-count expressions of *LRP1B*. **c**, Correlation of mean total counts (transcriptome sizes) between cell types in samples S1 and S2; CLTS normalization aims to align population-level transcriptome sizes toward unity (*y* = *x*). **d**, Relative adjustment ratios for CP10K-normalized expression (RNA) versus TF-IDF-normalized peak counts (ATAC) in matched L6 IT cells from multiome datasets (For visual clarity, the y-axis is truncated between 0 and 30; see Extended Data Fig. 2a for the unconstrained distribution.). **e–f**, Inter-sample Pearson correlation (*r*) of cell-type means for (**e**) transcriptome size and (**f**) chromatin accessibility (total peak counts) between any sample and the baseline. **g–h**, Multiomic integration analysis showing (**g**) Pearson correlation and (**h**) representative scatter plot of transcriptome size versus total peak counts within matched single-cell multiome pairs. All results are for the SEA-AD MTG data. **i, j,** Expression means of NEMGs in different types of cells of AD-dem, AD-non-dem and No-AD samples under CLTS (**i**) and TPM (**j**) normalization.

#### Scaling issues with CP10K normalization

Various methods have been developed to mitigate these issues, with CP10K (the default in Seurat^20^) being the most widely used. However, CP10K introduces significant scaling artifacts by forcing all cells to have a fixed 10,000 counts. This over-amplifies the expression profiles of cells with smaller transcriptome sizes (total counts) and suppresses those with large ones, causing biological misinterpretations (e.g., falsely indicating higher expression of the L5 IT marker *ST6GALNAC3* in microglia than in L5 ET neurons). Thus, despite reducing the inter-sample variance, CP10K distorts the cross-cell-type biological relationships (**Fig. 2a, CP10K**).

Besides distorting cross-cell-type comparisons, CP10K distorts expression patterns within cell types. As shown in **Fig. 2b**, CP10K significantly alters the expression profile of *LRP1B* in L6 IT cells from sample *S*1 compared to their matched raw counts as profiles of cells under the same sequencing depth were adjusted with different ratios. Our previous research^17^ demonstrated that other prominent methods, such as SCtransform^21^ and SCnorm^22^, suffer from similar scaling issues. To address this, we introduced CLTS, the first artifact-mitigated scRNA-seq normalization method^17^. CLTS removes cross-sample technical variation while preserving true biological signal (e.g., *ST6GALNAC3* elevation in L5 ET cells vs. microglia; **Fig. 1a, CLTS**). By leveraging the linear proportionality of mean total counts across samples, CLTS effectively harmonizes cellular profiles (**Fig. 2c**).

#### The core principle of CLTS

Because all cells within a given sequencing run share the same depth, the (measured) mean transcriptome sizes of all cell types maintain a constant ratio to their true biological sizes (i.e., their actual total RNA expression means). Consequently, the mean transcriptome sizes of various cell types should exhibit a strong linear correlation across samples, i.e., the mean transcriptome sizes of cell types in one sample are proportional, by a consistent ratio, to those of their corresponding cell types in another sample. For example, comparing samples *S*1 and *S*2 in **Fig. 2a**, the mean transcriptome sizes across cell types demonstrate a near-perfect linear association. Because *S*2 was sequenced at a higher depth, its observed values are approximately 2.1 times those of *S*1 (**Fig. 2c**). The core objective of CLTS is to leverage this linear relationship to harmonize the ratios between measured and true values across the entire dataset (**Fig. 2c**). By aligning these scaling factors to a common baseline and applying a uniform adjustment factor to all cells within the same sample (under the same sequencing depth), CLTS effectively mitigates sequencing depth artifacts while rigorously preserving the inherent biological variation between samples and cell types.

#### TF-IDF artifacts and multiomic inconsistencies

The snATAC-seq datasets are also highly sensitive to sequencing depth variations. Standard TF-IDF normalization^18, 19^, introduces scaling artifacts by forcing total peak counts per cell to a uniform value in the TF step. Consequently, TF-IDF distorts cellular accessibility profiles and disrupts the integration of chromatin accessibility and gene expression in snMultiome data. For example, in L6 IT neurons from a SEA-AD sample, ordering cells by expression scaling factors (CP10K) revealed near-stochastic TF-IDF scaling factors for matched accessibility profiles (**Fig. 2d; Extended Data Fig. 2a, b**). This artificial decoupling breaks the expected biological correspondence between modalities in single cells. To resolve this issue, we extended the CLTS framework to normalize snATAC-seq data.

#### Applicability of CLTS to snRNA-seq and snATAC-seq data

CLTS normalization scales target sample cell-type gene (or peak) count means to align with a reference baseline along the identity line y = x (**Fig. 2c**). This approach requires a strong linear correlation between the baseline and target cell-type means. In more than 99% of our snRNA-seq and snATAC-seq samples, baseline-to-target Pearson correlation coefficients (r) exceeded 0.9 (**Fig. 2e, f; Extended Data Fig. 2c, d**). Additionally, cell-type gene and peak count means correlated at r > 0.5 across all SEA-AD multiome samples (**Fig. 2g, h**). This confirms that global chromatin accessibility closely mirrors transcriptional output, validating CLTS application across both datasets.

#### Preserving biological signals with CLTS

CLTS effectively eliminates sequencing depth biases while preserving true biological signals. For example, NEMGs encode the subunits of the five OXPHOS complexes^23–26^ (**Table S1**). Because OXPHOS drives 90–95% of cellular ATP generation^27, 28^ and neurons consume 75–80% of the brain’s energy budget^29, 30^, NEMGs are expected to have higher expression in neurons than in non-neurons. Visium spatial transcriptomics confirmed this pattern, showing elevated NEMG expression in neuron-rich cortical regions compared to white matter (**Extended Data Fig. 3a**). CLTS-normalized pseudobulk RNA-seq profiles faithfully preserved this higher neuronal expression (**Fig. 2i; Extended Data Fig. 3b, c**). In contrast, conventional TPM normalization (the pseudobulk equivalent of CP10K) completely obliterates this signal, failing to show higher NEMG expression in neurons (**Fig. 2j; Extended Data Fig. 3d, e**). This benchmark clearly demonstrates that while the scaling artifacts of TPM mask essential biological realities, CLTS successfully uncovers and preserves them.

### Cell-type-specific dysregulation of nuclear-encoded oxidative phosphorylation complexes in AD dementia

To elucidate the molecular mechanisms underlying AD dementia, we generated cell-type-specific pseudobulk profiles from CLTS-normalized snRNA-seq data to compare AD-dem and AD-non-dem samples in the SEA-AD MTG region. Differentially expressed genes were heavily enriched for OXPHOS complexes I–V^23–26^ (**Table S1**), which comprise 13 mitochondrial-encoded subunits (MT-genes) and NEMGs. Examining NEMG expression revealed a striking cell-type-specific divergence: NEMGs were significantly upregulated in most non-neuronal cells but markedly downregulated in neurons (**Fig. 3a–e**).

**Fig. 3.**
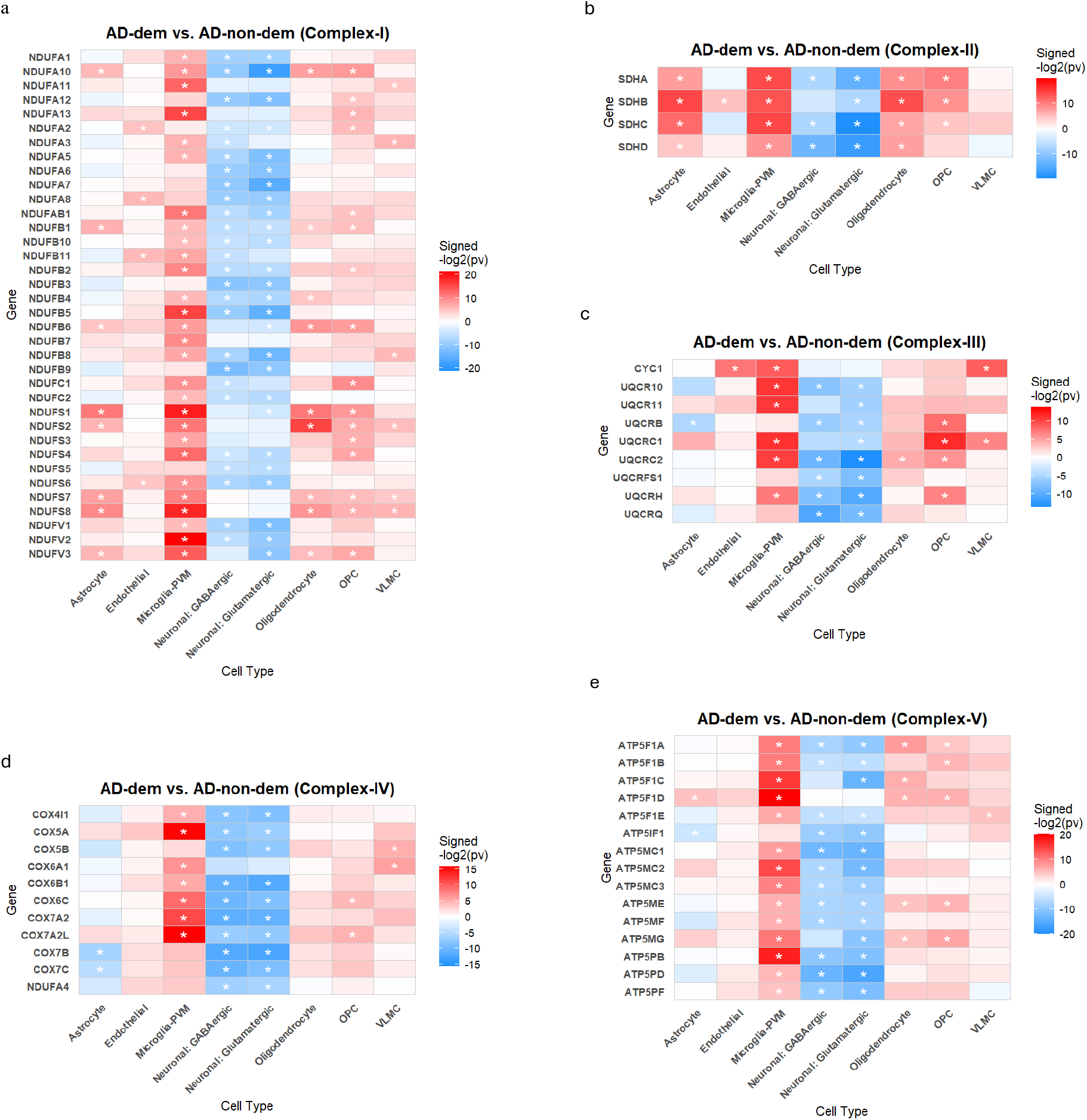
Cell-Type-Specific Dysregulation of OXPHOS Machinery in AD. **a–e**, Comparative expression of nuclear-encoded mitochondrial genes (NEMGs) across major cell types, categorized by OXPHOS enzyme complexes: (**a**) Complex I, (**b**) Complex II, (**c**) Complex III, (**d**) Complex IV, and (**e**) Complex V. Analysis contrasts AD-dem with AD-non-dem (resilient) samples. Asterisks indicate statistical significance (*p*_value < 0.05 via t-test).

#### Impaired energy homeostasis and NEMG downregulation in AD dementia

The human brain is exceptionally demanding, with neurons consuming approximately 20% of total bodily energy expenditure^31, 32^ and most of the brain’s energy budget^29, 30^. Although cells undergoing physiological stress in AD should theoretically exhibit heightened bioenergetic demands, transcriptomic profiling in the MTG reveals a striking paradox: NEMGs are profoundly suppressed in AD-dem samples rather than upregulated. This suppression likely disrupts OXPHOS complex assembly, resulting in ATP deficits that fail to meet the high neuronal energy demands^33, 34^. Extending our analysis to Brodmann area 9 (BA9)^4^ and the PFC^6^ confirmed this cell-type-specific pattern, non-neuronal NEMG upregulation alongside neuronal NEMG downregulation, though at a reduced magnitude compared to the MTG (**Extended Data Fig. 4**).

#### MT-gene elevation and mitonuclear decoupling mark the progression to AD dementia

All OXPHOS complexes, except complex II, encompass 13 MT-genes. We observed that the mean expression of MT-genes is significantly higher in neurons than in non-neurons within the MTG spatial transcriptomic dataset (**Extended Data Fig. 5a**) and CLTS-normalized pseudobulk data (**Extended Data Fig. 5b-d**). Like NEMGs, this cell-type-specific enrichment is lost under TPM normalization due to scaling issues (**Extended Data Fig. 5e-g**). Comparative analysis across datasets revealed that MT-genes were upregulated in AD-dem compared to AD-non-dem samples (**Extended Data Fig. 5h**), and in AD-non-dem compared to No-AD controls (**Extended Data Fig. 6a**) of major cell types. Crucially, the linear correlation between mean NEMG and MT-gene expression exhibits distinct patterns between neurons and non-neurons, offering a potential mechanism underlying the transition from pathological resilience to clinical dementia (**Extended Data Fig. 6b-i**). These specific findings will be further elaborated in the Discussion.

### Cell-type-specific suppression of mitochondrial biogenesis signaling in AD dementia neurons

In the preceding section, we demonstrated that the expression of NEMGs was significantly suppressed in neurons of AD-dem samples. This downregulation likely precipitated a neuronal bioenergetic crisis that caused AD dementia. Here, we studied the expression status of genes in the upstream mitochondrial biogenesis signaling.

#### Concomitant collapse of canonical and compensatory “bypass” biogenesis cascades in neurons

Under physiological energetic stress, neurons meet elevated ATP demands by activating a highly coordinated upstream and downstream network^35, 36^ that includes *PRKAA1*, *SIRT1*, *NRF1*, *NFE2L2* (*NRF2*), and *TFAM*. Within MTG glutamatergic neurons, while the expressions of these genes remained stable between AD-non-dem and No-AD controls, they were precipitously suppressed in the AD-dem samples (**Fig. 4a**).

**Fig. 4.**
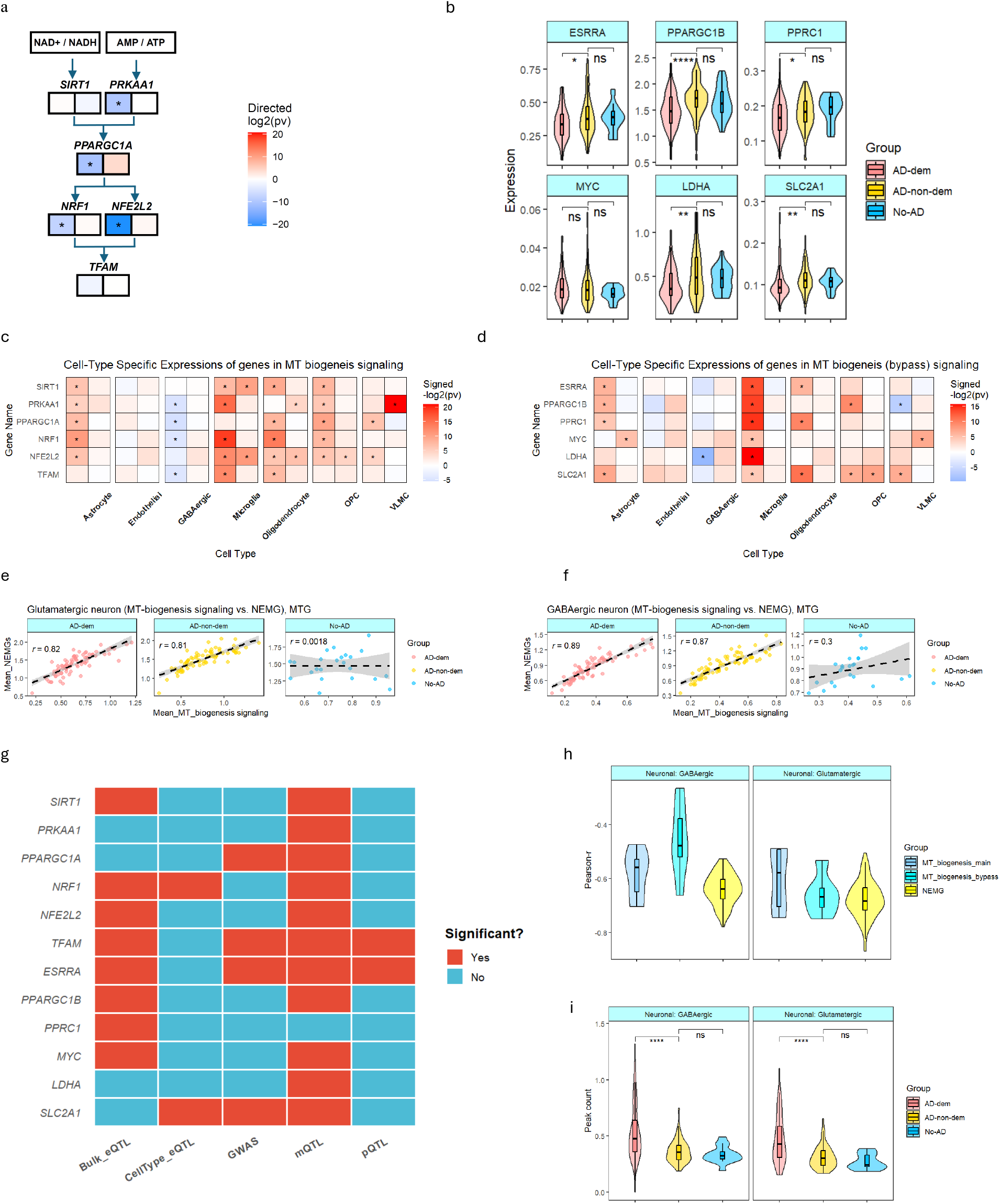
Metabolic and Transcriptional Drivers of Alzheimer’s Disease Progression. **a**, Differential expression of genes within the canonical mitochondrial biogenesis pathway in glutamatergic neurons. Heatmap cells are colored by **signed -*log*_2_(*p*-values)** to indicate both significance and direction of change (positive: up-regulated; negative: down-regulated). Comparisons are shown for AD with AD-dem vs. AD-non-dem (**left**) and AD-non-dem vs. No-AD (**right**). **b,** Expression profiles of “bypass” mitochondrial biogenesis signaling components across clinical patient groups in glutamatergic neurons. **c, d,** Cell-type-specific differential expression of canonical (**c**) and bypass (**d**) mitochondrial biogenesis genes across different cell populations. For each gene, the heatmap color scale representation is the same as those in (**a**). **e, f**, point-plot of expression mean of genes in mitochondrial biogenesis signaling and NEMGs for glutamatergic (**e**) and GABAergic (**f**) neurons. **g**, Status of identifying significant genetic variants in bulk eQTL, cell-type-specific eQTL, GWAS, pQTL, and mQTL analyses for genes involved in major and bypass mitochondrial biogenesis signaling pathways. **h**, Pearson correlation coefficient-r between chromatin accessibility (peak counts at chr12:120223689–120226293) and expression levels of genes in the major, bypass mitochondrial biogenesis signaling, and NEMGs in neuronal cells. **i**, Distribution of peak counts for the chr12:120223689–120226293 locus in neurons, stratified across AD-dem, AD-non-dem, and No-AD clinical samples.

This transcriptional silencing extends beyond the canonical axis. When the canonical PGC-1α ◊ NRF ◊ TFAM axis is compromised, neurons typically initiate alternative, compensatory “bypass” pathways to sustain bioenergetics and survival. Engagement of this bypass system conventionally relies on the upregulation of *ESRRA* (ERRα signaling)^37^, *PPARGC1B* and *PPRC1* (the PRC pathway)^38^, or *MYC* (the c-Myc pathway)^39^. In MTG glutamatergic neurons, however, most components of this bypass system were markedly downregulated in AD-dem samples while remaining stable across AD-non-dem and No-AD controls (**Fig. 4b**). GABAergic interneurons displayed a similar pattern of repression. In contrast, non-neuronal cells demonstrated stepwise upregulation of these genes across progressive AD stages (**Fig. 4c, d**).

To assess whether suppressing upstream regulators affects downstream targets, we evaluated the linear correlation between major biogenesis signaling and global NEMG expression levels. These correlations were remarkably robust across all major cell types in both AD-dem and AD-non-dem samples (**Fig. 4e, f; Extended Data Fig. 7a–f**), demonstrating that NEMG transcription remains actively tied to upstream signaling even in late-stage disease. Extending our profiling of major and bypass mitochondrial biogenesis signaling from the MTG to BA9^4^ and PFC^6^ revealed highly conserved transcriptional architectures across all three cortical regions (**Extended Data Fig. 7g, h**).

#### Divergent cell-type-specific biogenesis programs underlie AD dementia

In summary, neuronal NEMG suppression in AD-dem samples stems from a systematic, pan-pathway shut-off of canonical and compensatory biogenesis pathways, which could be driven by neuron-specific genetic and epigenetic changes. Conversely, non-neuronal cells upregulate these biogenesis signals and NEMGs in AD-dem relative to AD-non-dem stages. This divergence shows that non-neuronal functions remain intact into late-stage disease, enabling glia and vascular cells to meet the elevated bioenergetic demands needed to counteract neurodegeneration and maintain parenchymal tissue homeostasis.

### Identification of genetic drivers of AD dementia via multiomic xQTL and GWAS integration

To identify genetic variants driving the transition to AD dementia, we integrated genome-wide association study (GWAS) data with multiomic quantitative trait loci (xQTL) mapping. We systematically profiled single-nucleotide polymorphisms (SNPs) associated with genes involved in mitochondrial biogenesis signaling pathways and NEMGs using a multi-layered screening approach: bulk eQTL^40^, cell-type-specific eQTL^41^, pQTL^42^, mQTL^42^, and GWAS^43^ proximity scans. Candidates meeting these criteria were subsequently validated using Bayesian colocalization^44, 45^ and cis-Mendelian randomization (MR) analyses^46, 47^.

Our multiomic scans revealed widespread genetic regulation of these mitochondrial targets, with numerous variants significantly associated with gene expression, protein abundance, and/or DNA methylation levels, as well as proximal AD risk loci (**Fig. 4g, Extended Data Fig. 8**). Notably, the master mitochondrial regulators *TFAM* and *ESRRA* exhibited convergent evidence, harboring SNPs simultaneously associated with their mRNA expression, protein levels, and methylation profiles, along with adjacent AD-associated variants. This genetic architecture was even more pronounced among NEMGs; nearly all analyzed NEMGs possessed robust xQTL signals mapping to gene expression, protein levels, methylation state, or were paired with proximal AD-risk SNPs.

#### Prioritizing candidate variants with converging evidence

Multiomic integration prioritized *TFAM*, *ESRRA*, and *NRF1* as candidate regulators of mitochondrial biogenesis (**Supplementary Tables 2 and 3**; Supplementary Results). *TFAM* and *ESRRA* showed convergent eQTL, pQTL, and mQTL evidence, alongside suggestive regional AD GWAS associations. For *NRF1*, rs6949152 was reproducibly associated with lower expression in bulk DLPFC and excitatory neurons, although regional GWAS and colocalization support were limited in this case. These complementary patterns prioritize the three loci for functional follow-up without establishing causality.

Among the NEMGs, *NDUFS2* showed the strongest cross-platform convergence, combining a genome-wide significant DLPFC eQTL (rs35635728) and AD GWAS association (rs2070901) with moderate colocalization and complementary Mendelian randomization and methylation evidence (**Supplementary Tables 4 and 5**; Supplementary Results). Complex II genes *SDHA*-*SDHD* also carried strong bulk and cell-type eQTLs, but their regional GWAS signals did not reach genome-wide significance. Therefore, these loci are best viewed as regulatory candidates, whose relevance is reinforced by the consistent suppression of Complex II genes in AD-dem relative to AD-non-dem across the MTG and BA9.

*PPARGC1A* showed a predominantly epigenetic evidence profile, with a suggestive intralocus AD GWAS association and extensive brain mQTLs, but no significant eQTL, brain pQTL, or colocalization signal (**Supplementary Tables 2 and 3**; Supplementary Results). Together with its reduced expression in AD-dem neurons and the established role of PGC-1α in maintaining mitochondrial biogenesis^45, 46^, these data nominate *PPARGC1A* as an epigenetically sensitive regulator of neuronal vulnerability, although variant-level causality remains unresolved.

### Epigenetic regulation and identification of an upstream chromatin bottleneck

In this section, we leveraged the SEA-AD multiome dataset to investigate the epigenetic drivers of AD-associated transcriptional silencing across canonical biogenesis, alternative bypass pathways, and NEMGs. We mapped proximal and distal ATAC peaks correlated with paired gene expression (|Pearson r| ≥ 0.3) in glutamatergic and GABAergic neurons, identifying 93 candidate cis-regulatory elements in glutamatergic neurons and 12 in GABAergic neurons. Two peaks (chr8:142358634-142362266 and chr12:120223689-120226293) were conserved across both cell types (**Supplementary Tables 6–11**).

#### Epigenetic hyper accessibility at the PXN promoter drives mitochondrial network fragmentation in AD dementia

Genomic mapping revealed that the locus chr12:120223689-120226293 directly overlaps with the proximal *PXN* (Paxillin) promoter, adjacent to *COX6A1*, a core structural subunit of the OXPHOS complex IV (**Supplementary Table 12, 13**). Intriguingly, accessibility at this peak inversely correlates with expression across biogenesis networks, bypass pathways, and downstream NEMGs (**Fig. 4h**). Across disease stages, accessibility at this locus was elevated in AD-dem compared to AD-non-dem samples in both excitatory and inhibitory neurons, while AD-non-dem and non-AD samples showed similar levels (**Fig. 4i**).

Functional studies confirm that PXN hyperactivation or overexpression alters mitochondrial morphology, driving DRP1-mediated fission, network fragmentation, and metabolic oxidative stress^48–52^. Together, our multiomic integration suggests that AD-dem-specific epigenetic opening at the PXN promoter drives Paxillin overexpression. This focal adhesion scaffold disrupts mitochondrial networks and triggers retrograde mitonuclear signals, shutting down canonical and bypass biogenesis cascades in vulnerable neurons.

### Neurons exhibit metabolic shifts and ’energy crisis’ responses during AD progression

Downregulation of NEMGs and mitochondrial biogenesis pathways in AD-dem relative to AD-non-dem implicates a neuronal energy crisis in dementia (**Figs. 3, 4a–d**). To clarify the mechanisms of resilience and progression, we analyzed neuronal expression profiles across No-AD, AD-non-dem, and AD-dem stages.

#### Regulation of NEMGs and mitochondrial biogenesis in cognitive resilience and dementia

Across disease stages, NEMG expression in glutamatergic neurons subtly increased in AD-non-dem relative to No-AD, followed by marked suppression in AD-dem (**Fig. 5a**), a pattern maintained in GABAergic neurons (**Fig. 5b**). Importantly, these NEMG expression dynamics mirror the upstream mitochondrial biogenesis signaling (**Fig. 4a–f**). The initial upregulation of NEMGs in AD-non-dem suggests a compensatory response, wherein resilient neurons retain the capacity to sense metabolic stress and increase mitochondrial biogenesis to meet rising energy demands and stave off cognitive decline. However, in AD dementia, this homeostatic mechanism collapses: NEMG suppression impairs ATP production, ultimately driving cognitive failure.

**Fig. 5.**
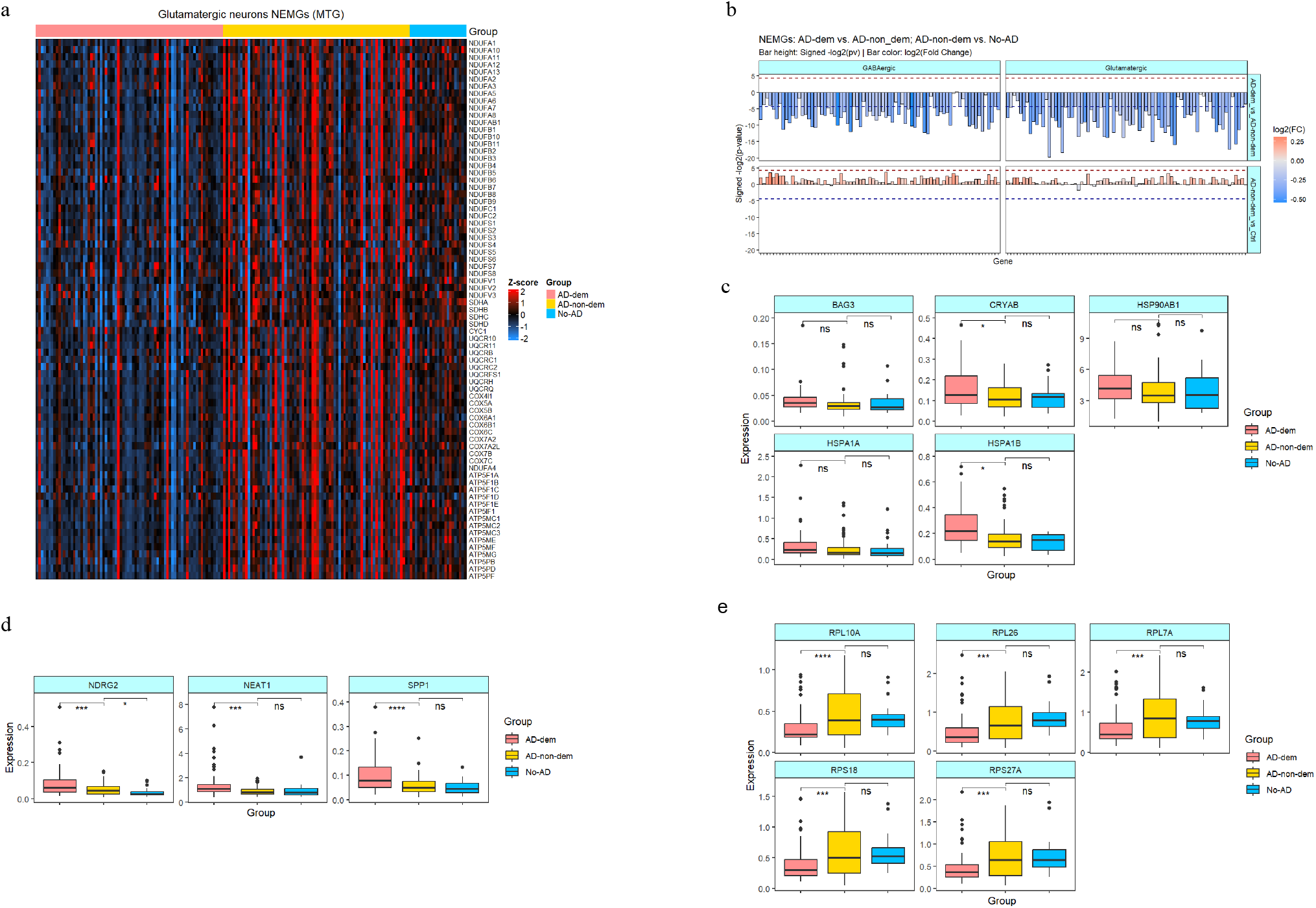
Metabolic and Proteostatic Dysregulation in Glutamatergic Neurons across AD Stages. **a**, Heatmap illustrating the expression profiles of NEMGs within glutamatergic populations across progressive stages of AD pathology. **b,** Comparative analysis of NEMG expression across clinical groups. Differential expression is displayed as signed -*log*_2_(*p*-values) and *log*_2_(fold-changes) for AD-dem vs. AD-non-dem (**top**) and AD-non-dem vs. No-AD (**bottom**) in both glutamatergic and GABAergic neurons. **c–e**, Expression dynamics within glutamatergic populations across AD progression, showing markers of (**c**) proteotoxic stress, (**d**) glial activation and the neuroinflammatory response, and (**e**) translational repression.

#### Downstream consequences of AD dementia samples: proteostatic stress and translational stagnation

Transcriptional decoupling disrupts energy homeostasis, leaving neurons unable to meet the high energetic demands of synaptic transmission. This “energy crisis” is supported by the coordinated induction of molecular chaperones, including *BAG3*, *CRYAB*, *HSP90AB1*, *HSPA1A*, and *HSPA1B*, in AD-dem neurons (**Fig. 5c**), indicating escalating proteotoxic stress^53–58^. Because protein folding and clearance are highly ATP-dependent, this heat-shock response may represent a final effort to preserve cellular integrity amid declining metabolic supply^59–61^.

Chronic bioenergetic distress is further indicated by upregulation of stress mediators *NEAT1*, *NDRG2*, and *SPP1* (**Fig. 5d**). Increased *NEAT1* and the AD-associated *NDRG2* suggest activation of adaptive stress and neuroprotective pathways under oxidative stress^62, 63^. Conversely, core ribosomal components, including *RPL7A*, *RPL10A*, *RPL26*, *RPS18*, and *RPS27A*, were broadly downregulated (**Fig. 5e**), consistent with reduced protein synthesis. Given the high energetic cost of translation, this translational suppression may conserve limited ATP resources^64^.

These metabolic and stress signatures were validated in the SEA-AD BA9^4^ region and PFC dataset^6^ (**Extended Data Fig. 9a-c**). Although the signals were most potent in the MTG, the patterns were consistent across cohorts. Collectively, these results suggest that while neurons in AD-non-dem samples successfully navigate an energy compensation mode, the transition to AD-dem is driven by a terminal energy crisis precipitated by the collapse of the nuclear-mitochondrial biogenic axis.

### Non-neuronal lineages exhibit sustained metabolic resilience and coordinated responses to AD pathology

Unlike neurons, which showed significant suppression of NEMGs and upstream mitochondrial biogenesis signaling in AD-dem samples, non-neuronal cells predominantly upregulated these genes (**Fig. 3**; **Fig. 4c, d; Extended Data Fig. 7g, h**). This divergence suggests that non-neuronal lineages face increased energy demands in advanced AD and activate mitochondrial biogenesis to meet these demands. Here, we examined how microglia/perivascular macrophages (Microglia-PVM) and astrocytes, key drivers of AD initiation, neuroinflammation, and progression, respond to distinct stages of pathology.

#### Sustained NEMG upregulation reflects functional and metabolic resilience in microglia-PVMs

In the SEA-AD MTG dataset, microglia-PVM NEMG expression peaks in AD-dem but remains comparable between AD-non-dem and No-AD controls (**Fig. 6a**). This suggests that microglia-PVM maintain metabolic robustness during advanced AD pathology, engaging biogenic signaling to meet higher energy demands. Except for astrocytes, other non-neuronal lineages display similar trajectories, reinforcing that these populations retain metabolic resilience across AD stages (**Fig. 6b**).

**Fig. 6.**
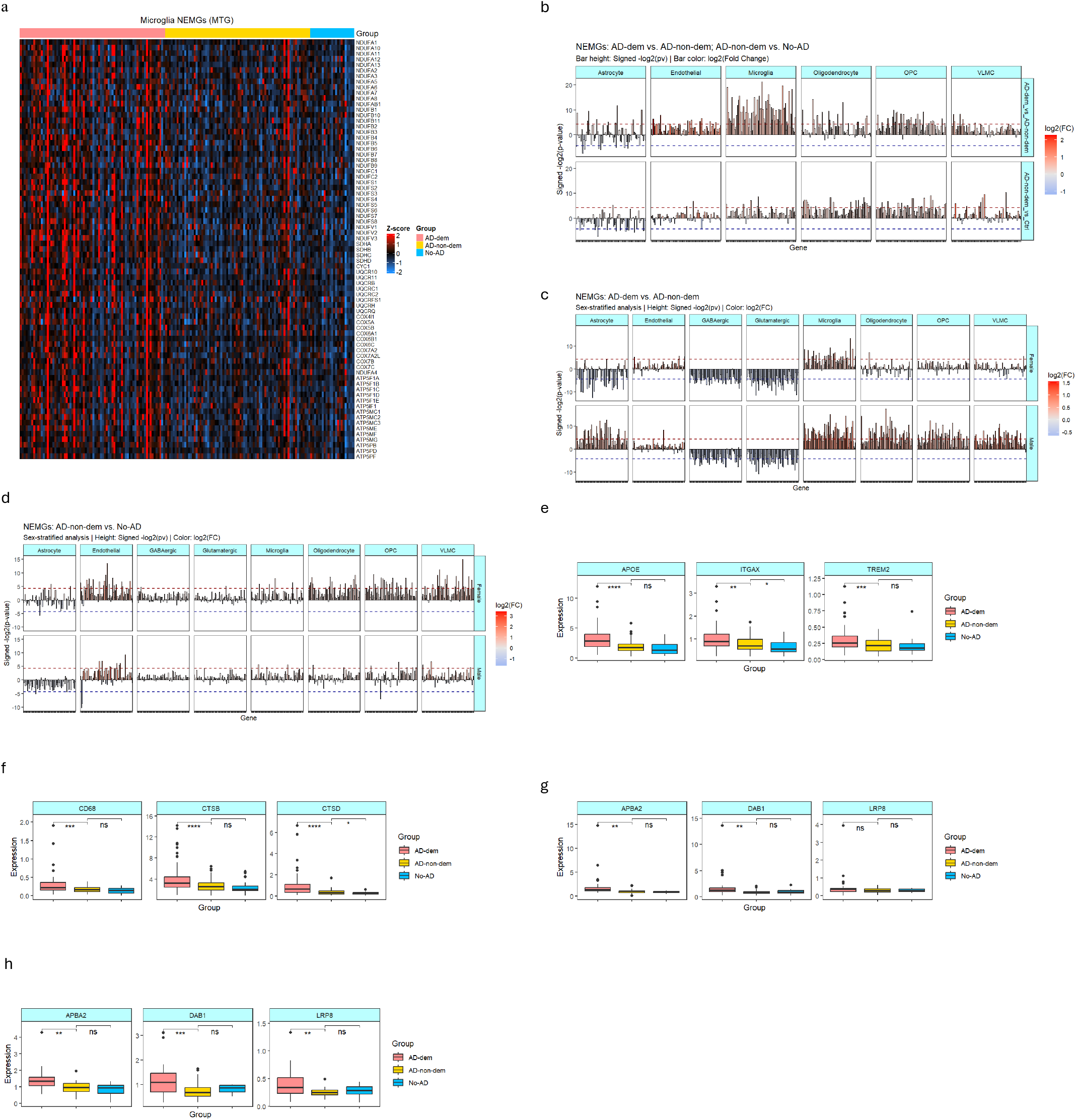
Metabolic Resilience and Adaptive Activation of Microglia-PVM across AD Stages. **a**, Heatmap illustrating the coordinated expression of NEMGs within Microglia-PVM across progressive stages of AD pathology. **b,** Comparative analysis of NEMG expression, displaying signed -*log*_2_(*p*-values) and *log*_2_(fold-changes) for AD-dem vs. AD-non-dem (**top**) and AD-non-dem vs. No-AD (**bottom**) across non-neuronal populations. **c, d,** Comparative analysis of NEMG expression in male and female samples, respectively, displaying signed -*log*_2_(*p*-values) and *log*_2_(fold-changes) for AD-dem vs. AD-non-dem (**c**) and AD-non-dem vs. No-AD (**d**). **e–h,** Boxplots depicting the progression of distinct functional signatures across AD pathology: (**e**) classic DAM markers, (**f**) “cleanup crew” genes involved in lysosomal degradation and phagocytosis, (**g**) astrocyte-mediated synaptic homeostatic support (**female samples**), and (**h**) astrocyte-mediated synaptic homeostatic support (**male samples**), where (**e, f**) for Microglia-PVM and (**g, h**) for Astrocyte.

#### Sex-stratified analyses revealed an opposite directional NEMG regulation unique to astrocytes

Sex-stratified analysis revealed a distinct metabolic divergence in astrocytes: in AD-dem, NEMGs were upregulated in males but downregulated in females, whereas other cell types like neurons remained directionally concordant between sexes (**Fig. 6c**). Astrocytes exhibited similar sex-dependent patterns when comparing AD-non-dem to No-AD controls (**Fig. 6d**). This divergence was conserved in BA9 astrocytes and present in oligodendrocytes across MTG and BA9, whereas neurons and microglia varied between sexes in response magnitude rather than direction (**Extended Data Fig. 9d, e**).

#### Coordinated microglial stress responses and DAM activation in AD

Beyond baseline metabolism, non-neuronal cells mounted an active response to AD stress. Microglia-PVM showed a progressive, stepwise upregulation of pathology-sensing genes (*APOE*, *TREM2*, *ITGAX*) across the diagnostic spectrum, significantly peaking in AD-dem relative to AD-non-dem (**Fig. 6e**). This elevated expression suggests enhanced plaque engagement and a transition toward a neuroprotective Disease-Associated Microglia (DAM) state^65–67^. Similarly, lysosomal and phagocytic markers (*CD68*, *CTSB*, *CTSD*) followed a matching upward trajectory (**Fig. 6f**), reflecting stepwise activation of microglial degradative machinery to process accumulating Aβ and tau burden^68–72^.

#### Preserved homeostatic and reelin signaling in astrocytes across sexes

A parallel compensatory astrocyte phenotype occurred across sexes: synaptic and homeostatic housekeeping genes (*APBA2*, *DAB1*, *LRP8*) were significantly upregulated in AD-dem compared to AD-non-dem, continuing an upward trajectory relative to controls (**Fig. 6g for females; Fig. 6h for males**). This elevation likely represents a direct response to neighboring neuronal distress and structural failure. Mechanistically, *APBA2* stabilizes APP processing to reduce toxic Aβ accumulation^73^, while *DAB1* and *LRP8* (*APOER2*) drive Reelin signaling, a pathway linked to modulating neuronal vulnerability in AD progression^13^.

#### Cross-regional conservation of non-neuronal adaptations

These non-neuronal responses were highly reproducible beyond the MTG dataset. In BA9^4^ and PFC^6^ snRNA-seq data, microglial activation markers (*APOE*, *TREM2*, *ITGAX*, *CD68*, *CTSB*, and *CTSD*) were consistently upregulated in AD-dem versus AD-non-dem microglia (**Extended Data Fig. 9f, g**). Likewise, *APBA2*, *DAB1*, and *RIMS1* remained elevated in AD-dem astrocytes across regions and sexes (**Extended Data Fig. 9h–j**). Together, these cross-regional findings demonstrate that non-neuronal lineages actively adapt via metabolic scaling and neuroprotective pathways to counteract the mounting neuronal energy stress.

### A unified model of cell type–specific metabolic divergence in Alzheimer’s disease progression

Our single-cell multiomic analyses revealed a stark functional dichotomy between neuronal and non-neuronal cells during the transition from the AD-non-dem to the AD-dem stages (**Extended Data Fig. 10**). While neurons undergo metabolic collapse marked by suppressed mitochondrial biogenesis signaling and NEMGs, glia, particularly microglia-PVM and astrocytes, mount a compensatory response with elevated metabolic machinery, DAM transition, and neuroprotective housekeeping functions. This lineage-specific divergence underscores a paradigm in which progressive neuronal energy failure parallels heightened bioenergetic engagement in the surrounding glia to manage the pathological burden.

## Discussion

### Overcoming scaling artifacts via CLTS normalization

Our work highlights a critical technical pitfall in bioinformatics: the scaling issue inherent in standard normalization methods like CP10K and TPM. We have shown before that under TPM normalization, the true metabolic dominance of neurons is masked, making MT-genes and NEMGs have higher expression in non-neurons (**Fig. 2j; Extended Data Fig. 3d, e; Extended Data Fig. 5e-g**). By utilizing CLTS, we avoid these scaling artifacts. CLTS allowed us to observe the true magnitude of NEMG and MT-gene expression in neurons, confirming that they are indeed the primary energy consumers (**Fig. 2i; Extended Data Fig. 3b, c; Extended Data Fig. 5b-d**). We propose that CLTS is a more robust standard for cross-cell-type comparisons, particularly when studying metabolic diseases like AD where total RNA output varies significantly between cell states.

Beyond confounding cross-cell-type comparisons, the scaling artifacts inherent to CP10K and TPM distort transcriptomic and epigenomic profiles within the same cell type and sample (**Fig. 2b, d; Extended Data Fig. 2a, b**). This phenomenon raises a fundamental methodological question: is it biologically or statistically justifiable to adjust individual cells from the same sample and sequencing library (i.e., with same sequencing depth) using vastly discrepant ratios? When standard methods like CP10K and TF-IDF are applied to snMultiome data, they scale matched multiomic profiles from the same individual cell using widely disparate ratios (**Fig. 2d; Extended Data Fig. 2a, b**). This artificial divergence systematically decouples the correlation between a cell’s transcriptional output and its chromatin accessibility. Consequently, these scaling artifacts compromise the accuracy of computational frameworks that rely on single-cell profiles, multiomic data integration (snRNA-seq and snATAC-seq), or joint genomic analyses (snRNA-seq and snWGS). By contrast, adopting CLTS for snRNA-seq and snATAC-seq normalization ensures a shared, biologically rigorous baseline. We anticipate that implementing CLTS will significantly improve the performance and fidelity of multiomic integration, joint modality modeling, and downstream computational pipelines across the single-cell field.

### Mitochondrial transcript dynamics as biomarkers for AD

A recurring debate in single-nucleus transcriptomics concerns the validity of MT-gene quantification, given that these transcripts primarily reside in the cytoplasm. However, our spatial transcriptomics (Visium) analysis demonstrates that neuron-enriched regions exhibit significantly higher MT-gene expression, directly corroborating the robust MT signals observed in multiple snRNA-seq data^4, 6^ (**Extended Data Fig. 5a-d**). These cell-type-specific patterns suggest that MT-gene expression in snRNA-seq data is a faithful biological reflection of neuronal metabolic demand rather than a technical artifact. This interpretation aligns with recent observations by Gabitto et al.,^4^ and Mathys et al.,^6^ further substantiating the role of MT-gene dynamics as informative biomarkers in Alzheimer’s disease pathology.

### MT–NEMG Transcriptional Decoupling as a Marker of Dementia

Our most striking finding was a shifting linear correlation between MT-genes and NEMGs across disease states in neurons. While No-AD controls showed a moderate positive correlation, AD-non-dem samples shifted to a moderate negative correlation (**Extended Data Fig. 6b, c**). We interpret this shift as an “energy compensation mode”: as NEMG expression drops, neurons upregulate MT-genes to maintain ATP production, preserving cognitive function despite the underlying AD pathology.

In AD-dem samples, this co-expression correlation was markedly attenuated, signaling a breakdown in mito-nuclear coordination (“transcriptional decoupling”) (**Extended Data Fig. 6b, c**). In this state, neurons fail to boost MT-gene expression against progressive NEMG suppression, compromising OXPHOS complex assembly and precipitating an irreversible energy crisis that drives dementia developments.

Conversely, non-neuronal cells maintained strong positive correlations across all stages (**Extended Data Fig. 6d–i**), suggesting that their mito-nuclear coordination remains functionally intact even in late-stage disease.

Physiologically, master biogenesis cascades should tightly synchronize MT-gene and NEMG transcription, a relationship preserved in non-neurons but disrupted in AD neurons. This mitonuclear transcriptomic uncoupling demonstrates that NEMG suppression enables autonomous drivers of MT-gene expression, which is aligned with mouse studies in which ATP depletion (13–60%) selectively stimulated MT-gene transcription without altering NEMG expression^74^. Taken together, these data strongly suggest that under severe bioenergetic deficit, neurons deploy an emergency retrograde signaling mechanism to upregulate MT-genes independently of canonical, nuclear-driven biogenesis pathways.

## Online content

Supplementary information, including the Methods and Discussion sections, tables, and figures, can be found in the Supplementary section.

## Acknowledgements

This research was partially supported by National Institutes of Health grants awarded to Z.Z. (U01AG079847 and R01LM012806). We thank the technical support from Cancer Prevention and Research Institute of Texas (CPRIT RP240610). C.C. is a CPRIT Postdoctoral Fellow in the Biomedical Informatics, Genomics and Translational Cancer Research Training Program (BIG-TCR) funded by Cancer Prevention & Research Institute of Texas (CPRIT RP210045 and RP260802). The funders had no role in the study design, data collection and analysis, decision to publish, or preparation of the manuscript.

## Author contributions

**S.L.** and **Z.Z.** conceived the project. **S.L.** developed and implemented CLTS_mtx, performed data analyses and model evaluations, and drafted the manuscript. **N.E.** performed the GWAS and xQTL analyses. **C.C.** assisted with data collection, preprocessing, Peak site gene mapping, figure design, and drafting the manuscript. **C.H.T.** and **Z.Z.** contributed to the discussion of results. **C.H.T.** contributed figure design. All authors reviewed and commented on the manuscript at all stages.

## Data availability

The single-nucleus RNA sequencing (snRNA-seq) and single-nucleus ATAC sequencing (snATAC-seq) datasets analyzed in this study are publicly available through the Seattle Alzheimer’s Disease Brain Cell Atlas (SEA-AD) portal (https://sea-ad-single-cell-profiling.s3.amazonaws.com/index.html) and the Synapse database under accession ID syn51123521.

## Code availability

The source codes and Tutorials are freely available online at GitHub: https://github.com/bsml320/CLTS_mtx and zenodo repository https://zenodo.org/records/21708359.

---

**Extended Data Fig. 1.**
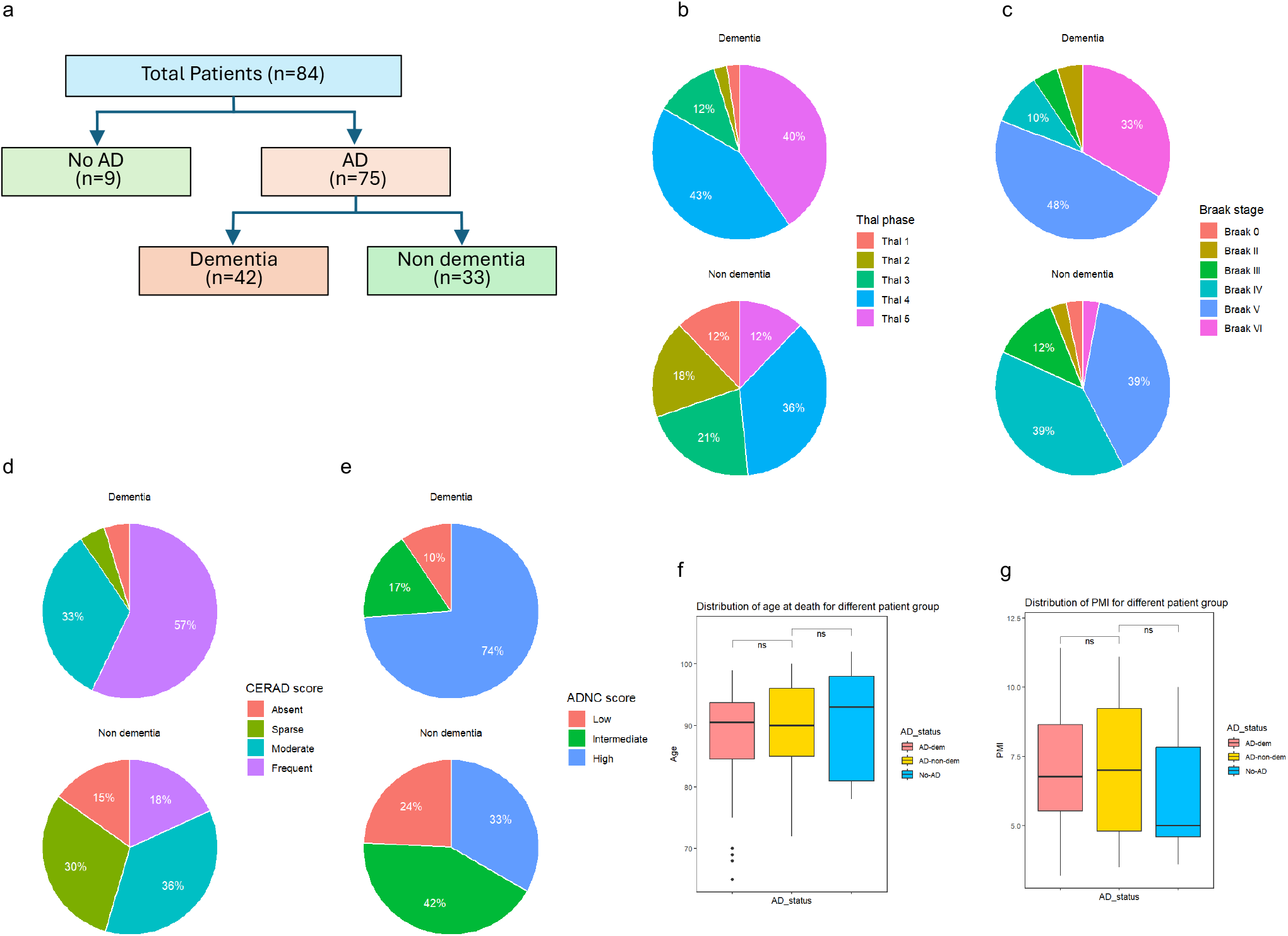
Clinical and Neuropathological Characterization of the SEA-AD Cohort. **a**, Stratification of donors by clinical sample classification. **b–e**, Comparison of AD-related neuropathology scores, including (**b**) Thal phase (amyloid-β plaque distribution), (**c**) Braak stage (neurofibrillary tangle progression), (**d**) CERAD score (neuritic plaque density), and (**e**) ADNC score (AD neuropathological change), stratified by AD Dementia and AD No-Dementia status. **f, g,** Comparison of the distribution of age (**f**) and post-mortem interval (PMI) (**g**) of patients in different AD pathology groups.

**Extended Data Fig. 2.**
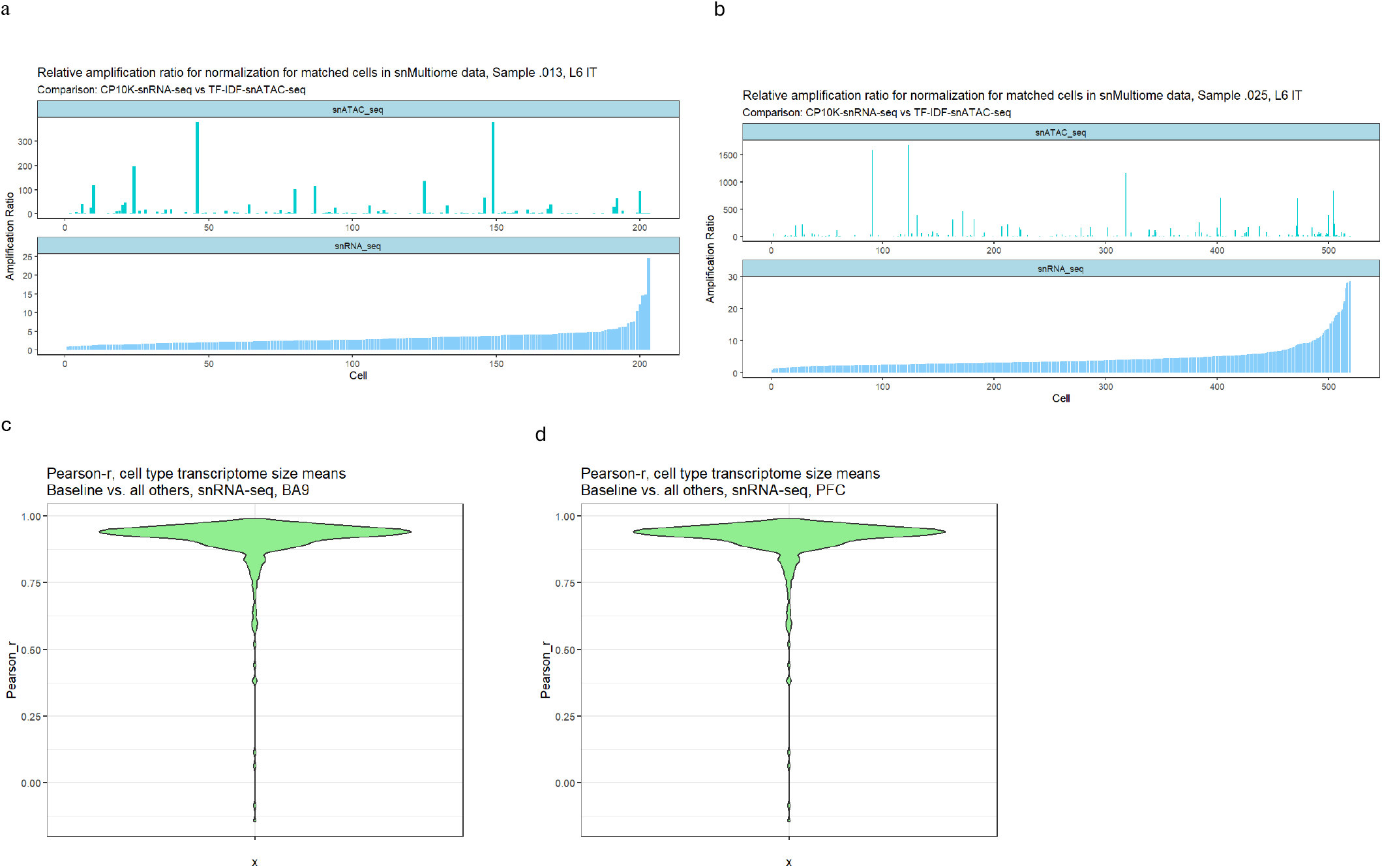
Benchmarking CLTS normalization across samples. a, b, Relative adjustment ratios for CP10K-normalized expression (RNA) versus TF-IDF-normalized peak counts (ATAC) in matched L6 IT cells from multiome datasets for sample .013 (**a**) and sample .025 (**b**). **c, d**, Inter-sample Pearson correlation (*r*) of cell-type means for transcriptome size between any samples to the baseline for BA9 (**c**) and PFC (**d**) regions.

**Extended Data Fig. 3.**
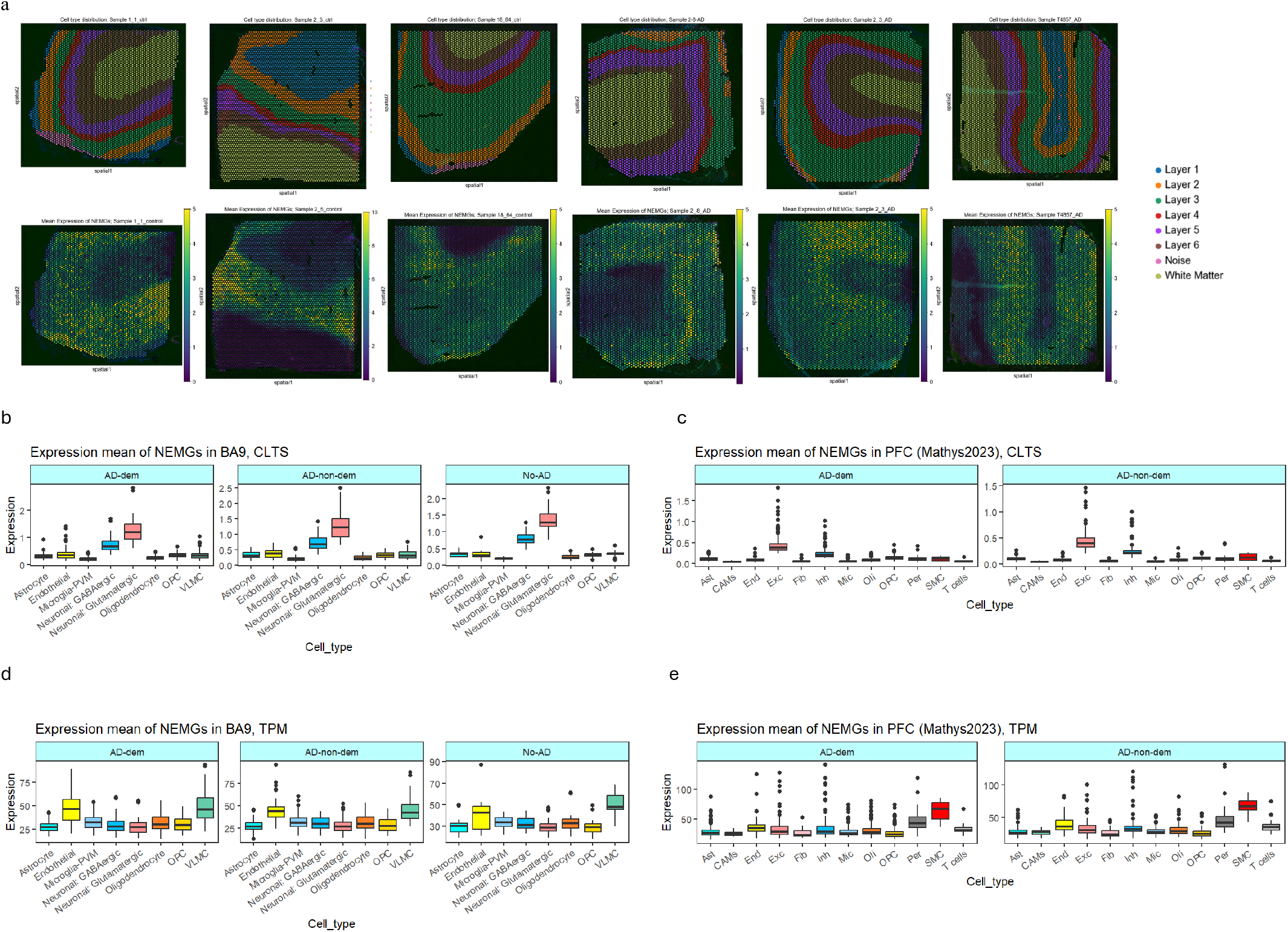
Comparison of NEMG expression between neurons and non-neuronal cells across Visium spatial transcriptomics and snRNA-seq datasets. **a**, Mean expression of NEMGs across Visium spots from six human brain MTG tissue samples (top: anatomical layers; bottom: expression means of NEMGs). **b**, **c**, Mean expression of NEMGs across distinct cell types from AD-dem, AD-non-dem, and No-AD samples under CLTS normalization, evaluated in the SEA-AD BA9 dataset (**b**) and the Mathys et al. PFC dataset (**c**). **d, e**, Mean expression of NEMGs across distinct cell types from AD-dem, AD-non-dem, and No-AD samples under TPM normalization, evaluated in the SEA-AD BA9 (**d**), and Mathys et al. PFC (**e**) datasets.

**Extended Data Fig. 4.**
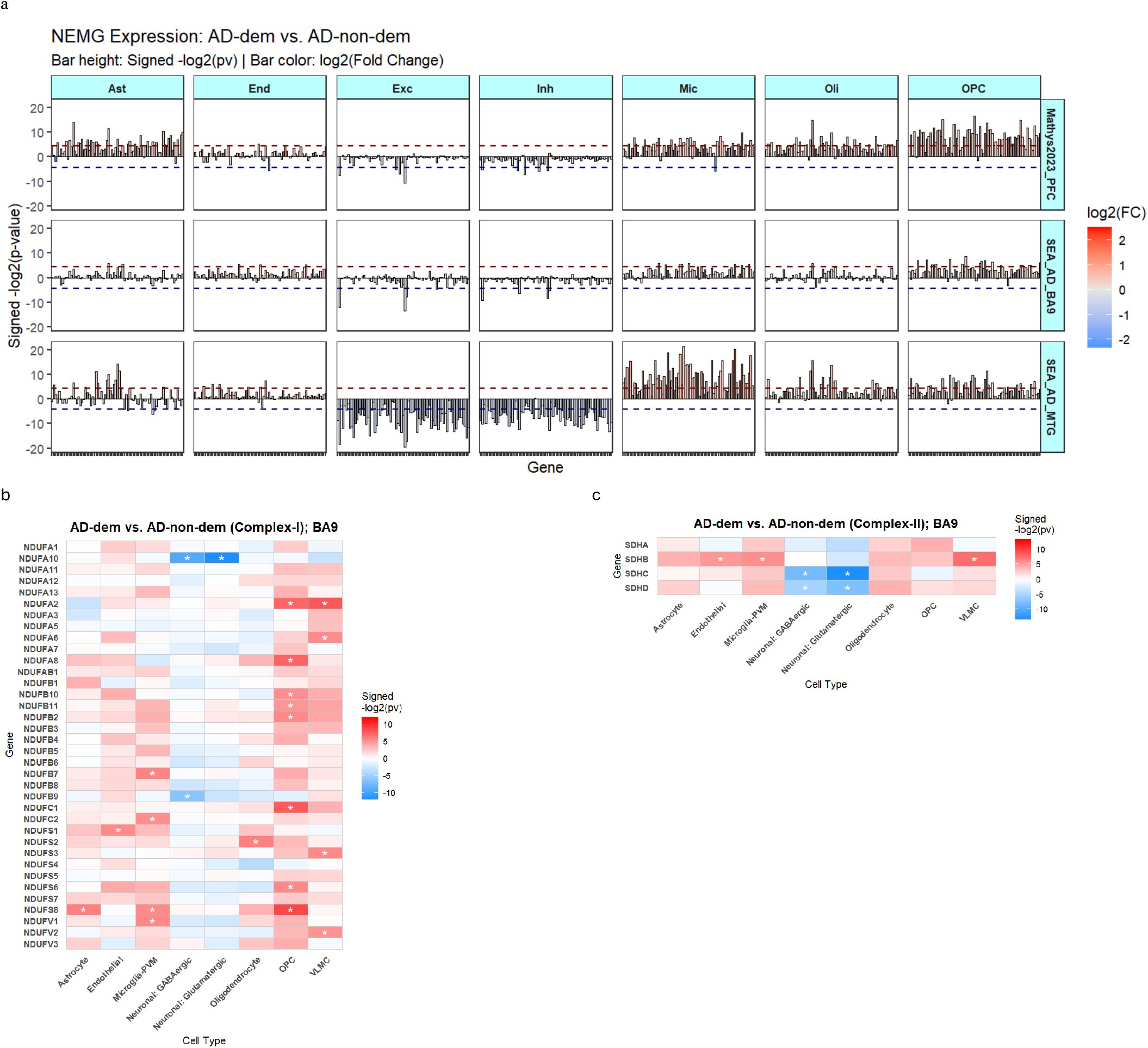
Comparison of NEMG expression between neurons and non-neuronal cells across snRNA-seq datasets. **a**, Differential expression analysis of NEMGs comparing AD-dem versus AD-non-dem conditions. Plots display signed -*log*2(p-values) and *log*2(fold-changes) across the SEA-AD MTG (**bottom**), SEA-AD BA9 (**middle**), and Mathys et al. PFC (**top**) datasets. **b, c,** Comparative expression of nuclear-encoded mitochondrial genes (NEMGs) across major cell types of BA9 region, categorized by OXPHOS enzyme complexes: (**b**) Complex I, (**c**) Complex II. Asterisks indicate statistical significance (*p*_value < 0.05 via t-test).

**Extended Data Fig. 5.**
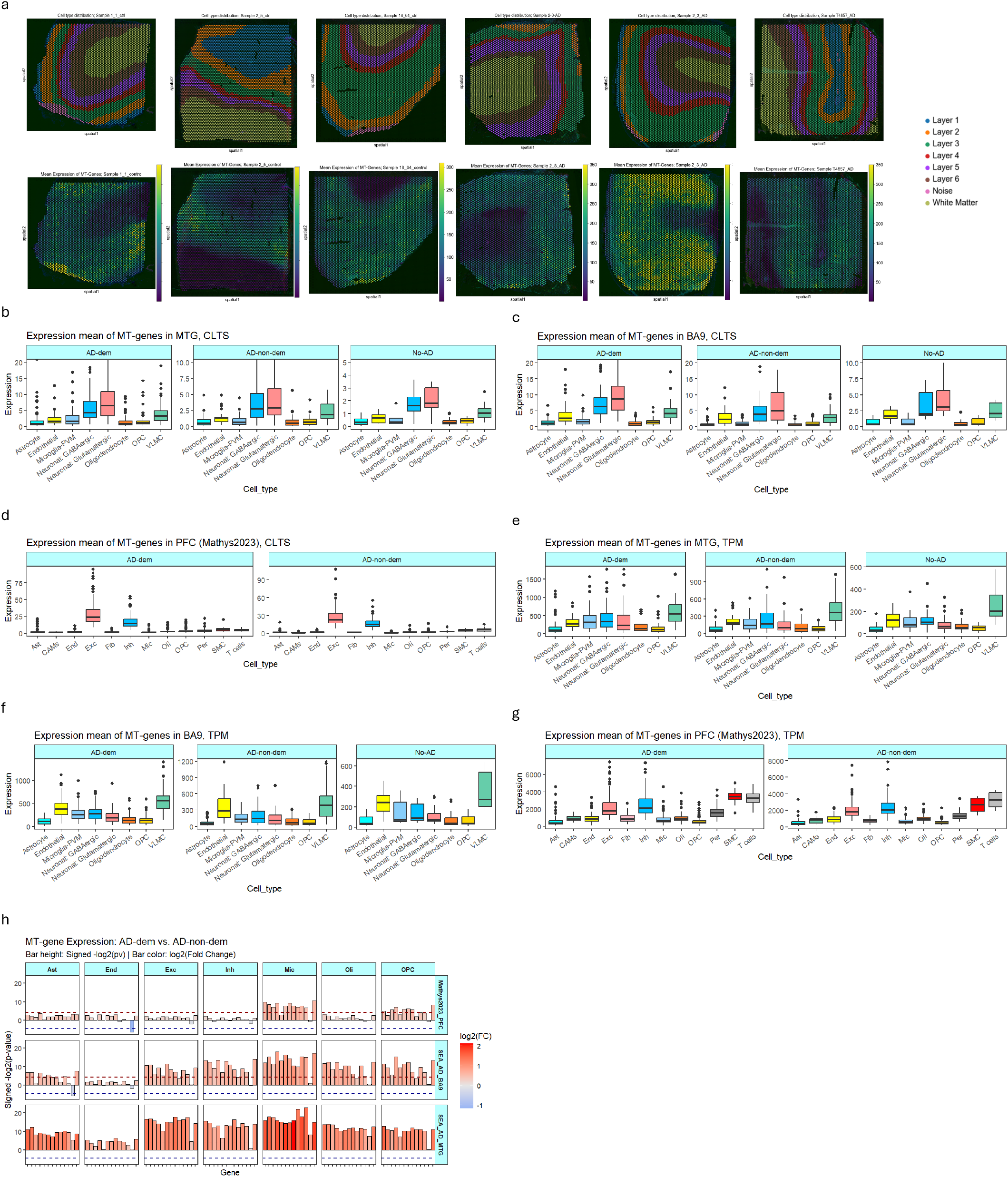
Comparison of MT-gene expression between neurons and non-neuronal cells across Visium spatial transcriptomics and snRNA-seq datasets. **a**, Mean expression of MT-genes across Visium spots from six human brain MTG tissue samples (top: anatomical layers; bottom: expression means of MT-genes). **b-d,** Mean expression of MT-genes across distinct cell types from AD-dem, AD-non-dem, and No-AD samples under CLTS normalization, evaluated in the SEA-AD MTG dataset (**b**), BA9 (**c**) and the Mathys et al. PFC dataset (**d**)**. e–g,** Mean expression of MT-genes across distinct cell types from AD-dem, AD-non-dem, and No-AD samples under TPM normalization, evaluated in the SEA-AD MTG (**e**), SEA-AD BA9 (**f**), and Mathys et al. PFC (**g**) datasets. **h,** Differential expression analysis of MT-gene comparing AD-dem versus AD-non-dem conditions. Plots display signed -*log*_2_(p-values) and *log*_2_(fold-changes) across the SEA-AD MTG (**bottom**), SEA-AD BA9 (**middle**), and Mathys et al. PFC (**top**) datasets.

**Extended Data Fig. 6.**
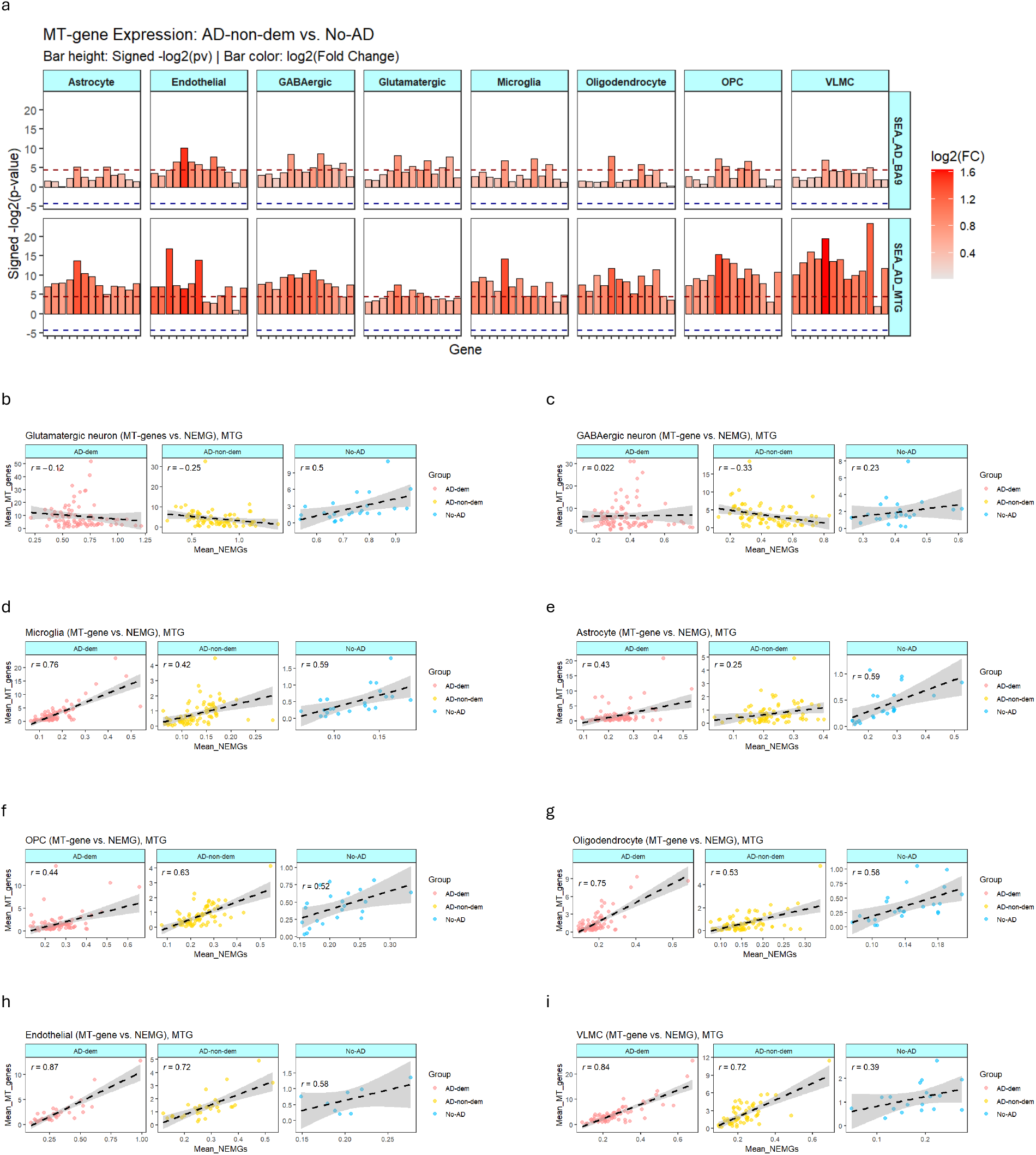
Correlation and differential expression analysis of nuclear-encoded mitochondrial genes (NEMGs) versus mitochondrial-DNA-encoded genes (MT-genes). **a**, Differential expression analysis of MT-genes comparing AD-non-dem versus No-AD conditions. Plots display signed -*log*_2_(p-values) and *log*_2_(fold-changes) across the SEA-AD MTG (**bottom**), SEA-AD BA9 (**top**) datasets. **b–i**, Scatter plots showing the relationship between mean expression of NEMGs and MT-genes across distinct cell types, including glutamatergic neurons (**b**), GABAergic neurons (**c**), Microglia-PVM (**d**), astrocytes (**e**), OPCs (**f**), oligodendrocytes (**g**), endothelial cells (**h**), and VLMCs (**i**).

**Extended Data Fig. 7.**
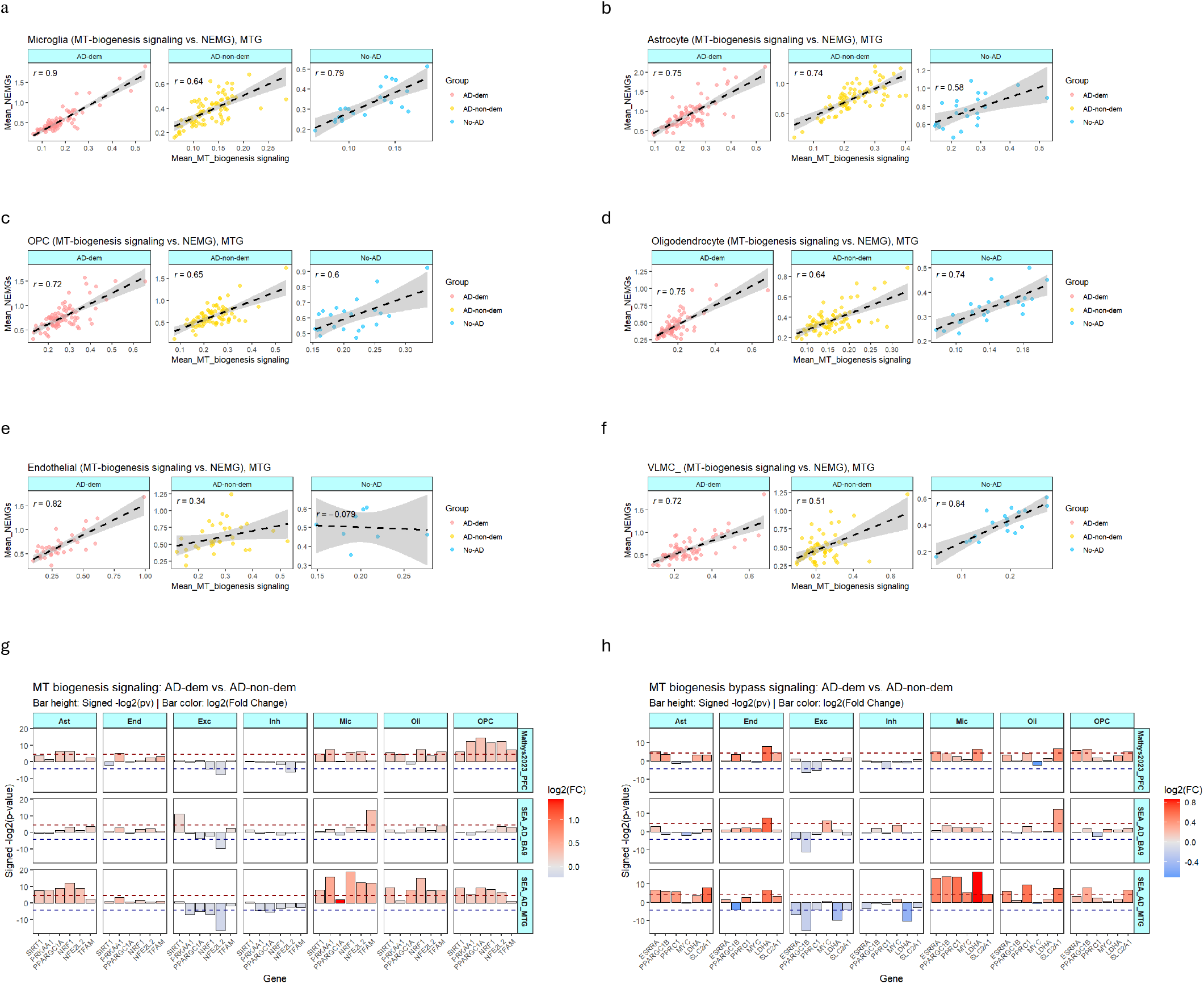
Correlation of nuclear-encoded mitochondrial genes (NEMGs) versus mitochondrial biogenesis signaling. **a–f**, Scatter plots showing the correlation between mean expression of NEMGs and MT-genes across non-neuronal cell types, including Microglia-PVM (**a**), astrocytes (**b**), OPCs (**c**), oligodendrocytes (**d**), endothelial cells (**e**), and VLMCs (**f**). **g, h**, Differential expression analysis of canonical mitochondrial biogenesis signaling (**g**) and bypass signaling (**h**) across distinct cell types and datasets. Plots display signed -*log*2(*p*-values) and *log*2(fold-changes) for the comparison of AD-dem versus AD-non-dem.

**Extended Data Fig. 8.**
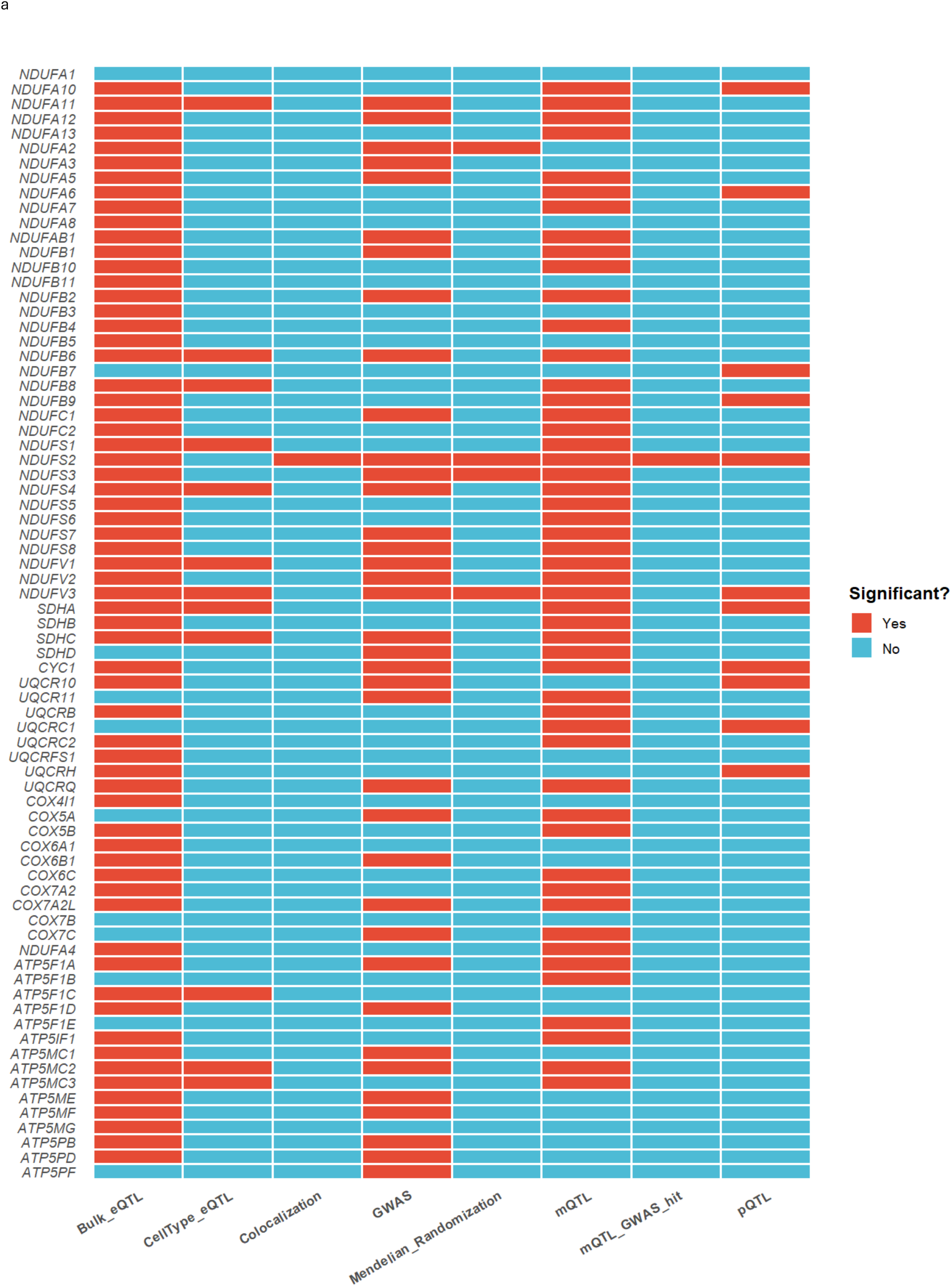
SNPs associating with the expression, protein abundance, and methylation of NEMGs. **a**, Landscape of NEMGs with SNPs achieving statistical significance across bulk tissue and cell-type-specific eQTLs, disease GWAS, pQTLs, and mQTLs, colocalization and mendelian randomization.

**Extended Data Fig. 9.**
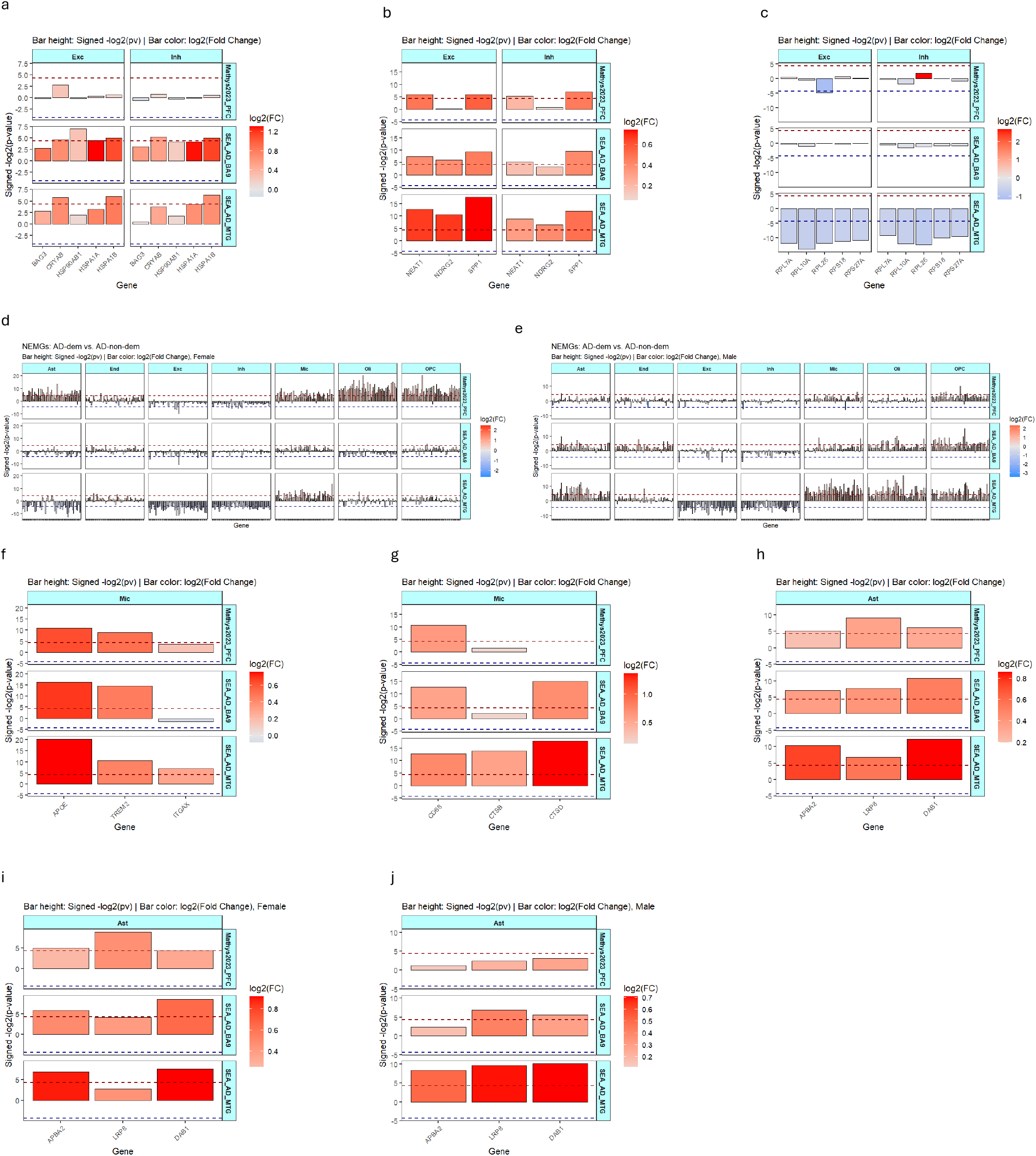
Transcriptional changes in functional response pathways to neuronal energetic crisis. **a–c**, Differential expression analysis of downstream functional response cascades to metabolic stress across specific cell types and clinical cohorts. Plots display signed *log*2(*p*-values) and *log*2(fold-changes) for the AD-dem versus AD-non-dem comparison, highlighting:(**a**) proteotoxic stress signatures in glutamatergic neurons, (**b**) glial activation and neuroinflammatory response pathways in glutamatergic neurons, (**c**) translational repression markers in glutamatergic neurons. **d**, **e**, Differential expression analysis of NEMGs across distinct cell types and datasets comparing AD-dem versus AD-non-dem in female (**a**) and male (**b**) cohorts. Plots display signed *log*2(*p*-values) and *log*2(fold-changes). **f–j**, Differential expression analysis of downstream functional response cascades to metabolic stress across specific cell types and clinical cohorts. Plots display signed *log*2(*p*-values) and *log*2(fold-changes) for the AD-dem versus AD-non-dem comparison, highlighting, (**f**) canonical DAM markers in microglia, (**g**) lysosomal degradation and phagocytic (“cleanup crew”) genes in microglia, (**h**) astrocyte-mediated synaptic homeostatic support across all combined samples, (**i**) astrocyte-mediated synaptic homeostatic support restricted to female samples, and (**j**) astrocyte-mediated synaptic homeostatic support restricted to male samples.

**Extended Data Fig. 10:**
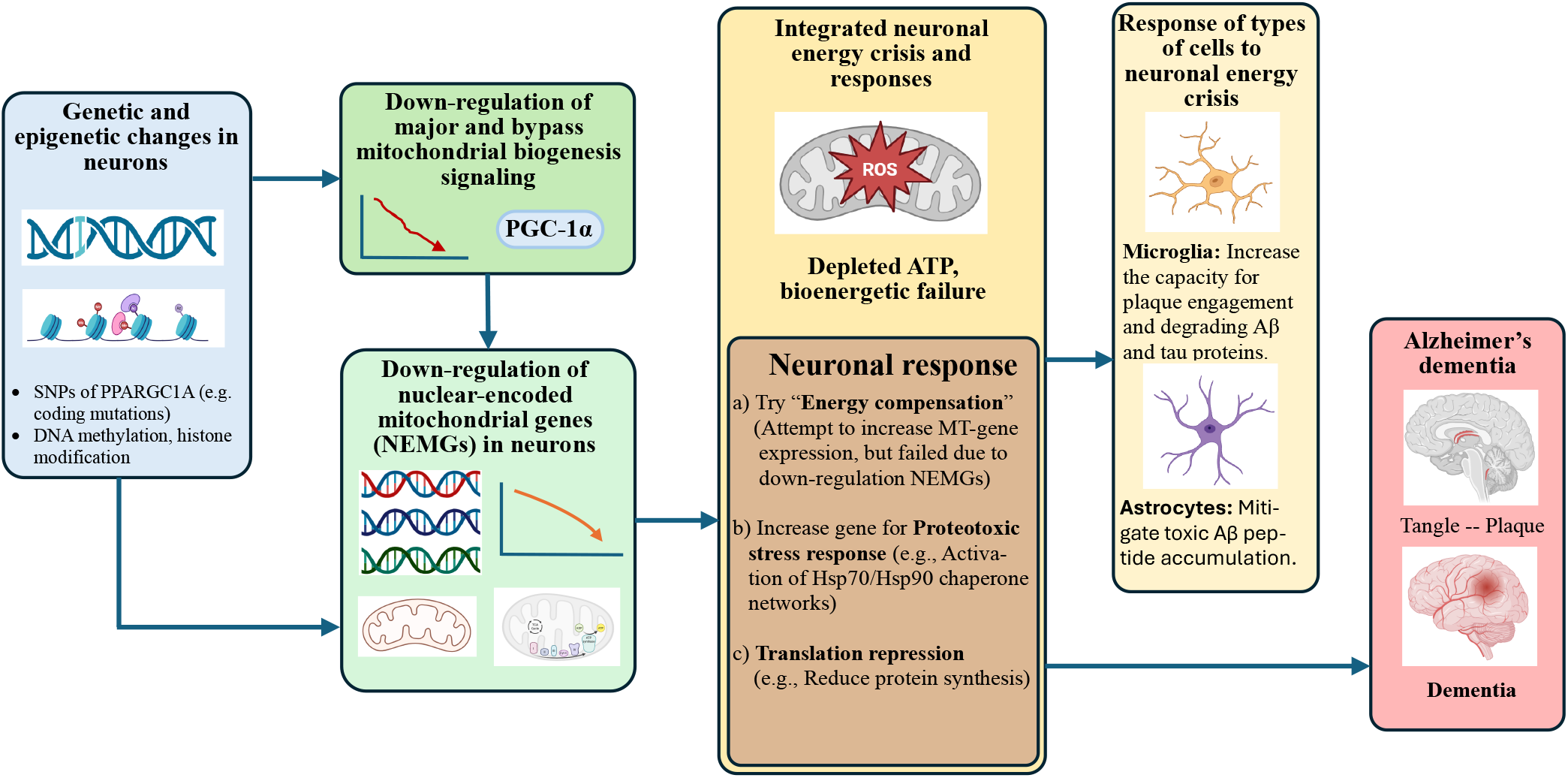
Proposed disease mechanism driving AD dementia. Neuronal genetic and epigenetic alterations downregulate master regulators of mitochondrial biogenesis signaling. This suppression impairs NEMGs across all five complexes of the OXPHOS chain, triggering an energy crisis in neurons. This metabolic dysfunction compromises neuronal integrity and drives the transition to AD dementia. In response to this neuronal energy crisis, surrounding glial populations—including microglia and astrocytes—undergo compensatory or reactive functional shifts in AD dementia brains. (Icons created with BioRender.com).

