## Supplementary material for "Single-Nucleus Profiling Reveals Neuronal Metabolic Collapse During the Transition to Alzheimer’s Disease Dementia": Supp.

### Supplementary Results

#### Multiomic evidence supporting candidate variants in mitochondrial regulatory pathways

**Detailed evidence for *TFAM*, *ESRRA*, and *NRF1*.** Integration of bulk eQTL, AD GWAS, brain pQTL, and brain mQTL data revealed distinct but complementary evidence profiles for these three mitochondrial regulators (**Supplementary Tables 2 and 3**). For *TFAM*, rs58326258 was a strong bulk eQTL in CommonMind DLPFC ($P = 4.71 \times 10^{-20}$; $\beta = -0.227$), whereas the strongest regional AD GWAS association was rs1819658 ($P = 1.80 \times 10^{-6}$; 232 kb from the gene). The locus also showed a brain pQTL association (Q00059; $P = 2.00 \times 10^{-15}$) and 465 significant brain mQTL associations across seven CpGs, led by cg13123656 in the parahippocampal gyrus. For *ESRRA*, rs473578 was associated with expression in GTEx cerebellum ($P = 3.69 \times 10^{-5}$; $\beta = -1.51$), and the leading regional GWAS variant, rs11231727, reached $P = 3.59 \times 10^{-5}$ at 60.7 kb from the gene. This locus was further supported by a brain pQTL (P11474; $P = 1.82 \times 10^{-7}$) and 1,150 significant mQTL associations across five CpGs, with cg27160968 as the strongest DLPFC signal. *NRF1* showed cross-resolution eQTL concordance: rs6949152 was associated with lower expression in both ROSMAP DLPFC ($P = 1.21 \times 10^{-12}$; $\beta = -0.527$) and excitatory neurons ($P = 1.52 \times 10^{-7}$). The regional GWAS evidence was weaker (rs549019941; $P = 4.68 \times 10^{-4}$; 34 kb from the gene) and did not support colocalization (PP.H4 = 0.0125), but the locus contained 819 significant mQTL associations spanning 12 CpGs, led by cg24892351 in DLPFC.

**Detailed evidence for *NDUFS2* and Complex II genes.** Among the NEMGs, *NDUFS2* showed the clearest cross-platform convergence (**Supplementary Tables 4 and 5**). The bulk DLPFC eQTL rs35635728 reached genome-wide significance ($P = 2.12 \times 10^{-8}$), and the locus contained a genome-wide significant AD GWAS association at rs2070901 ($P = 1.05 \times 10^{-8}$; 0.545 kb from the gene). Bulk eQTL and GWAS signals showed moderate colocalization (PP.H4 = 0.5105). Mendelian randomization implicated rs1136224 ($\beta = 0.147$; $P = 1.12 \times 10^{-4}$), although its quality-control flag (standard-error ratio > 3) warrants caution. Independent support came from a brain pQTL (O75306; $P = 7.40 \times 10^{-8}$) and 734 significant mQTL associations across 18 CpGs, including methylation-GWAS overlap at rs4575098 (GWAS $P = 1.33 \times 10^{-8}$). Lead bulk eQTLs for Complex II mapped to rs7718951-*SDHA* ($P = 4.83 \times 10^{-15}$), rs2076601-*SDHB* ($P = 2.15 \times 10^{-8}$), rs558623569-*SDHC* ($P = 4.30 \times 10^{-23}$), and rs4026118-*SDHD* ($P = 5.92 \times 10^{-5}$). Excitatory-neuron eQTLs further supported rs2977280-*SDHB* ($P = 4.60 \times 10^{-4}$) and rs10157990-*SDHC* ($P = 1.34 \times 10^{-9}$). These four genes carried 1,958, 2,294, 1,504, and 1,108 significant mQTL associations, respectively, and *SDHA* showed a strong brain pQTL ($P = 1.47 \times 10^{-21}$). Their regional GWAS minima did not reach genome-wide significance ($P = 1.30 \times 10^{-5}$ to $2.78 \times 10^{-4}$), supporting their interpretation as regulatory candidates rather than established AD-risk alleles.

**Detailed evidence for *PPARGC1A*.** No significant bulk or cell-type eQTL, brain pQTL, or variant-level colocalization signal was detected for *PPARGC1A* in the analyzed datasets (**Supplementary Tables 2 and 3**). Among 8,283 SNPs evaluated around the locus, rs191019573 produced the strongest AD GWAS association ($P = 2.15 \times 10^{-6}$) and mapped within the gene. The methylation signal was broader: 1,909 significant brain mQTL associations were identified across 31 CpGs, led by cg25659893, although no individual mQTL overlapped an AD GWAS hit. This pattern supports epigenetic regulation of *PPARGC1A* but does not resolve a causal variant. Its biological relevance is reinforced by lower *PPARGC1A* expression in glutamatergic and GABAergic neurons from AD-dem donors and by prior experimental evidence that loss of PGC-1α suppresses NEMG expression[1], whereas its overexpression increases mitochondrial biogenic capacity[2]. Independent studies have linked rs8192678 to reduced PGC-1α expression and metabolic insufficiency[3-5].

### Methods

#### CLTS_mtx: Scalable Single-Nucleus Multi-Omics Normalization

Although our previously developed method, CLTS, effective for generating robust reference profiles for bulk RNA-seq cell-type deconvolution, was not designed to scale efficiently to massive datasets. Because its original scope was limited to targeted sub-samples, processing high-throughput single-nucleus RNA-seq (snRNA-seq) and ATAC-seq (snATAC-seq) data resulted in severe time and memory bottlenecks. To address these constraints, we re-engineered the framework into CLTS_mtx. By natively processing sparse matrix structures (**.mtx format**) and optimizing core algorithmic routines, CLTS_mtx dramatically reduces computational overhead, enabling rapid, memory-efficient normalization of large-scale multiomics datasets.

The core theoretical foundation of CLTS_mtx remains identical to CLTS, resting on three principal hypotheses:

1. **Estimating Biological Ground Truth**: Data points obtained from snRNA-seq represent discrete measurements of underlying biological ground truth. Specifically, the mean transcriptome size (total count) of a given cell type serves as a robust estimator of the average total RNA molecules expressed by that population.
2. **Uniform Intra-sample Technical Bias**: For all cells within a single sequencing library or a sample, the measured values maintain a constant ratio relative to their true biological values. This implies that technical factors, such as capture efficiency and amplification bias, apply uniformly across all cell types within the same sample.
3. **Variable Inter-sample Proportionality**: Sequencing depth discrepancies manifest as significant variations in this ratio between different samples.

Under our hypothesis, the mean transcriptome sizes of all cell types should show a strong linear correlation across samples, as the mean sizes in one sample are expected to be proportional to their counterparts in another. For example, when comparing samples S1 and S2, the mean cell-type transcriptome sizes demonstrate a near-perfect linear relationship. Because S2 was sequenced to a higher depth, its observed values are uniformly scaled by a factor of approximately 2.1 relative to S1 (**Fig. 2c**). The primary objective of CLTS_mtx is to leverage this linear relationship to harmonize the ratio of expression profles across the entire dataset (**Fig. 2c**). Specifically, CLTS_mtx applies a uniform adjustment factor to all cells within a given sample, reflecting the physical reality of snRNA-seq and snATAC-seq protocols, where cells in the same library share a common sequencing depth. The open-source CLTS_mtx package, along with comprehensive documentation, is publicly available on GitHub at https://github.com/bsml320/CLTS_mtx.

**Generation of Cell-Subtype-Specific Pseudobulk Profiles**

To generate cell-subtype-specific pseudobulk profiles for each sample, we aggregated the CLTS-normalized RNA sequencing (snRNA-seq) and single-nucleus ATAC sequencing (snATAC-seq) datasets. For a given sample and cell type with at least 10 cells, the pseudobulk transcriptomic profile was constructed by calculating the mean normalized count for each gene across all constituent cells. An identical averaging strategy was applied to the snATAC-seq peak matrix to generate corresponding cell-type-specific pseudobulk chromatin accessibility profiles.

**Downstream Analysis of Major Cell-Type Transcriptomes**

Differential expression analysis was performed independently for each major cell type. Differentially expressed genes (DEGs) were identified by executing two parallel pairwise comparisons using the cell-type-specific pseudobulk matrices: AD dementia (AD-dem) versus AD no-dementia (AD-nodem), and AD-nodem versus non-AD controls. Downstream functional profiling focused on the transcriptional dynamics of the oxidative phosphorylation (OXPHOS) machinery. Specifically, we systematically evaluated the expression trajectories of nuclear-encoded mitochondrial genes (NEMGs) and the 13 mitochondrial-encoded genes (MT-genes) structural units across the five multi-subunit OXPHOS complexes. Finally, to map the regulatory architecture underlying these metabolic shifts, we analyzed the expression profiles of upstream transcriptional pathways governing mitochondrial biogenesis, alongside gene networks managing downstream cellular responses to neuronal energy crises.

**Identification of Genetic Drivers via GWAS and eQTL Integration.**

This analysis applies the full 5-step genetic evidence pipeline — eQTL Catalogue v2 Scan (Bulk Brain), Bryois 2022 Cell-type eQTL Scan, GWAS Proximity Scan, Colocalization (coloc.abf), cis-Mendelian Randomization, pQTL and mQTL Scan (FunGen-xQTL Portal)— to the genes of interest, including NEMGs in the five OXPHOS complexes and genes in the mitochondrial biogenesis signal. This analysis used 19-dataset tabix scan, Bryois 2022 cell-type eQTLs, and GWAS proximity scan. This extended analysis uses the identical methodology applied to the genes of interest.

**Gene Coordinates**. GRCh38 coordinates fetched via Ensembl REST API (/lookup/symbol/homo_sapiens/{gene}) for all 27 genes. All confirmed on correct chromosomes and cross-verified against eQTL scan chromosome assignments.

**eQTL Catalogue v2 Scan (Bulk Brain).** Tabix-based queries to EBI FTP (ftp://ftp.ebi.ac.uk/pub/databases/spot/eQTL/sumstats/{study_id}/{dataset_id}/{dataset_id}.all.tsv.gz) for 19 brain gene-expression (GE) datasets across 6 studies. Cis-window: gene midpoint ± 500 kb (= 1 Mb total). For each gene × dataset, all SNPs in the window were extracted and filtered to the target Ensembl gene ID. Datasets used:

| Dataset ID | Study | Tissue | n |
|---|---|---|---|
| QTD000075 | CommonMind | DLPFC | 452 |
| QTD000434 | ROSMAP | DLPFC | 494 |
| QTD000051 | BrainSeq | DLPFC | 209 |
| QTD000166 | GTEx | Cerebellum | 241 |
| QTD000161 | GTEx | Cerebellar hemisphere | 209 |
| QTD000176 | GTEx | DLPFC | 197 |

| Dataset ID | Study | Tissue | n |
|---|---|---|---|
| QTD000579 | Walker 2019 | Neocortex | 212 |
| QTD000146–206 | GTEx | 10 additional brain regions | 84–205 |
| QTD000041–046 | Braineac2 | Putamen, Substantia nigra | 84/93 |

Significance thresholds: genome-wide significant (GW-sig) $p < 5\times10^{-8}$; suggestive $p < 1\times10^{-5}$.

**Bryois 2022 Cell-type eQTL Scan.** All 104 files (8 cell types × 13 chromosomes) downloaded from Zenodo record 7276971. Cell types: Astrocytes, Endothelial cells, Excitatory neurons, Inhibitory neurons, Microglia, OPCs/COPs, Oligodendrocytes, Pericytes. Files are plain gzip (not bgzipped); scanned by decompressing and filtering by Ensembl gene ID. SNP genomic positions resolved via the accompanying snp_pos.txt file (GRCh38 coordinates, 5.34M SNPs).

**Note**: Bryois files lack standard error (SE) columns; colocalization is not possible for Bryois-only signals.

**GWAS Proximity Scan.** The full Bellenguez 2022 GWAS file (21,101,114 SNPs) was streamed from EBI FTP in a single pass. For each of the genes of interest, all SNPs within a ±500 kb window around the gene midpoint were extracted (172,035 total matched SNPs; 955–10,781 per gene). For each gene: (1) minimum p-value in the window and its lead SNP, (2) nearest genome-wide significant hit ($p < 5\times10^{-8}$) and its distance from the gene body. Distance measured from SNP position to nearest gene boundary (0 if within gene body).

**Colocalization (coloc.abf).** For each GW-sig gene ($p < 5\times10^{-8}$ in eQTL Catalogue v2), full regional eQTL summary statistics were extracted from the per-gene tabix scan files. Bellenguez 2022 GWAS regional slices were extracted from the streaming scan (±500 kb windows). SNPs matched by genomic position; INDELs excluded from coloc (type ≠ 'SNP'); duplicates removed. coloc.abf run with default priors ($p1 = p2 = 1\times10^{-4}$, $p12 = 1\times10^{-5}$). eQTL dataset: type = "quant", N from dataset metadata. GWAS dataset: type = "cc", N = 788,989, s = 0.1411 (111,326 cases / 788,989 total). Posterior probabilities reported: PP.H0 (no association), PP.H1 (eQTL only), PP.H2 (GWAS only), PP.H3 (distinct causal variants), PP.H4 (shared causal variant = colocalization).

**Cis-Mendelian Randomization.** Wald ratio cis-MR was performed for all genes with genome-wide significant bulk brain eQTLs ($p < 5\times10^{-8}$) for which GWAS data were available. *SDHA* was excluded (instrument rs7718951 at chr5:297,024 is subtelomeric and absent from Bellenguez 2022 GWAS — same limitation as in the Set 2 analysis). Instruments were required to be SNPs (INDELs excluded). Allele alignment between eQTL alt allele and GWAS effect allele was verified for all 9 instruments using allele data fetched directly from the Bellenguez 2022 GWAS file; GWAS beta was sign-flipped where eQTL alt = GWAS other allele. GWAS imputation quality assessed by SE ratio (observed/expected SE at instrument MAF, n =

788,989); instruments with SE ratio > 10 excluded. MR SE computed by delta method: MR SE = |GWAS SE / eQTL β|. P-values from two-sided z-test.

**pQTL Scan (FunGen-xQTL Portal).** Brain proteomics pQTL data queried from the FunGen-xQTL NIAGADS portal for each of the 75 genes. Data source: ROSMAP DLPFC (n=328–543) and MSBB PHG (n=328) TMT mass spectrometry proteomics. For each gene, the portal was queried for cis-pQTL associations (FDR-significant, within ±1 Mb of gene TSS). Genes with at least one FDR-significant cis-pQTL were classified as pQTL-positive. Colocalization with Bellenguez 2022 GWAS was assessed using Wakefield approximate Bayes factors (ABF) with Z-scores from the portal and GWAS beta/SE. PP.H4 ≥ 0.5 was the threshold for strong colocalization.

**mQTL Scan (FunGen-xQTL Portal).** Brain methylation mQTL data queried from the FunGen-xQTL NIAGADS portal for each gene. Data source: ROSMAP DLPFC, Knight-ADRC, and MSBB brain methylation (450K array, n=543). For each gene, all FDR-significant cis-mQTL associations (SNP–CpG pairs within ±1 Mb of gene TSS) were retrieved. GWAS variant overlap was assessed by matching mQTL SNP IDs against Bellenguez 2022 GWAS hits ($p < 5\times10^{-8}$) using rsID string matching. **Limitation:** variants encoded as chr:pos:ref:alt strings (rather than rsIDs) in either dataset are silently excluded from this matching step, meaning the overlap test may under-detect true overlaps. Formal coloc.abf was not run because the portal provides only FDR-significant variants (~5% of the cis window), which systematically underestimates PP.H4; variant overlap is reported instead.

**Identification of Epigenetic Drivers via Single-Nucleus Multiomic Analysis**

To identify potential epigenetic drivers of metabolic dysfunction, we performed a peak-to-gene correlation analysis independently within the glutamatergic and GABAergic neuronal populations. For each lineage, chromatin accessibility at specific peak sites was correlated against the mean expression profiles of core genes involved in primary mitochondrial biogenesis signaling, bypass mitochondrial biogenesis pathways, and nuclear-encoded mitochondrial genes (NEMGs). Candidate regulatory elements were restricted to peak sites exhibiting a robust linear relationship, defined by a Pearson correlation coefficient of $|r| \geq 0.3$. Conserved regulatory features were then isolated by identifying overlapping peak sites between the glutamatergic and GABAergic datasets. Finally, these conserved peaks were mapped to potential target genes using a 500kb cis-regulatory window centered on the transcriptional start site (TSS).
